# Single-cell analysis of West Nile virus infection in brain organoids identified a serine synthesis enzyme as a determinant of viral neural tropism and a druggable antiviral target

**DOI:** 10.64898/2026.10.23.761475

**Authors:** Michael Leitner, Lachlan De Hayr, Ziyao Zhao, Freyja E Stam, Jia Xiang Justin Chan, Washington Romero, Nicole Quigg, Abigail L Cox, Nathan Godde, Daniel J Rawle, Mohammed Shaker, Ernst Wolvetang, Alexander A Khromykh, Andrii Slonchak

## Abstract

The cellular and molecular determinants of human neuroinvasive West Nile virus (WNV) disease are currently poorly defined. Previous studies of WNV neural infection relied on rodent models and immortalised cell lines that diverge from human brain tissue. Herein, we established a reproducible human cerebral organoid model of WNV infection and profiled it by virus-inclusive single-cell RNA sequencing (scRNA-Seq) to obtain an understanding of WNV-host interactions in the context of human neural tissue.

We found that WNV showed preferential tropism to progenitor cells, immature neurons, and astrocytes, while the mature neurons were largely resistant to infection. Astrocytes were identified as the principal source of interferon (IFN) and the main hub of IFN-signalling, whereas mature neurons expressed little viral RNA-sensing machinery or STAT1 and responded weakly to infection and to IFN. The differential gene expression analysis between WNV-permissive and nonpermissive cells demonstrated that virus-susceptible cells had high baseline expression of the serine synthesis and one-carbon metabolism genes *PHGDH*, *SHMT2*, and *MTHFD2*, which were not induced by infection. RNA-interference and pharmacological inhibition of phosphoglycerate dehydrogenase (PHGDH) restricted WNV replication, confirming the requirement of this gene for WNV infection. In addition, the FDA-approved drug disulfiram, known to inhibit PHGDH, also reduced viral replication in neural progenitors and cerebral organoids.

Overall, these results provide the first comprehensive single-cell-level characterisation of WNV neural infection in human neural tissue. They identify *de novo* serine synthesis as a metabolic dependency of WNV in human neural cells that can be targeted as a host-directed antiviral therapy.

## INTRODUCTION

West Nile virus (WNV) is a mosquito-borne positive strand RNA virus in the Orthoflavivirus genus of the *Flaviviridae* family. It represents the most geographically widespread cause of arboviral encephalitis worldwide (*1*). In nature, the virus is maintained in an enzootic cycle between *Culex* mosquitoes and avian amplifying hosts, while humans and horses represent incidental dead-end hosts that cannot sustain virus transmission (*2*). Since its introduction into North America in 1999, WNV has become endemic across the Americas and various virus strains continue to circulate in Europe, Africa, the Middle East, and Australia (*3*). Most human infections are asymptomatic or cause a self-limiting febrile illness, but in some cases, they can progress to West Nile neuroinvasive disease, which presents as meningitis, encephalitis, or acute flaccid paralysis. Advanced age and immunosuppression are considered to be the risk factors of West Nile neuroinvasive disease (*4*). WNV is also an important veterinary pathogen that causes severe and frequently fatal encephalomyelitis in horses (*5*). Despite the substantial burden of WNV infection, there is no licensed human vaccine or approved antiviral therapy and clinical management remains entirely supportive (*6*).

WNV isolates segregate into two principal genetic lineages. Lineage 1 has a worldwide distribution and includes the strains responsible for most major epidemics of neuroinvasive disease such as the North American NY99 strain (*7*). Lineage 2 is historically confined to sub-Saharan Africa and was long regarded as comparatively avirulent. However, in recent decades it emerged across Europe, where it now causes recurrent outbreaks of severe neuroinvasive disease (*8*). In Australia, WNV is represented by the endemic Kunjin strain (WNV_KUN_), a lineage 1 variant historically associated with sporadic and mild disease in humans and horses (*9*). However, this epidemiological picture changed in 2011, when an unprecedented outbreak of equine encephalitis across southeastern Australia caused neurological disease in close to a thousand horses with high mortality (*5*). The outbreak was attributed to the emergence of a more neurovirulent WNV_KUN_ variant, designated WNV_NSW2011_ (*5*). Relative to historical Australian Kunjin isolates, WNV_NSW2011_ is also substantially more neuropathogenic in rodent models, which has been linked to several mutations in the viral genome (*10*). A case of human encephalitis has also been reported for WNV_NSW2011_ infection (*11*). These features define WNV_NSW2011_ as a veterinary- and clinically-relevant WNV strain with encephalitic potential (*5, 12*).

Despite the global burden of WNV neurological disease, the cellular and molecular mechanisms of WNV encephalitis remain poorly understood. Following neuroinvasion, WNV infects cells of the central nervous system (CNS), yet the cell-intrinsic determinants of viral tropism to CNS cells, and the cell-type-specific innate immune responses within the human brain are unclear (*13*). Past studies of WNV neurovirulence and neuropathogenesis relied primarily on mouse models (*14–17*) and cultured human and murine cells (*18–20*). However, rodents differ from humans in antiviral immunity, including interferon responses and restriction-factor repertoires (*21, 22*). Immortalised human cell lines and two-dimensional neural cultures lack the cellular diversity, and maturation of human brain tissue, and frequently carry transformation-associated alterations in innate immune signalling (*23*). Primary human brain tissue and derived cultures are informative (*24*), although scarce and may not be amenable to controlled infection. Human induced pluripotent stem cell (iPSC)-derived cerebral organoids address many of these constraints as they represent self-organising, three-dimensional structures that recapitulate key aspects of human brain tissue architecture and contain the major neural cell types, including neural progenitors, radial glia, astrocytes, and neurons (*25*). Cerebral organoids have been used extensively to model the neuropathogenesis of multiple viruses including Zika virus (*26–31*) and SARS-CoV-2 (*32*), and are now considered as the most relevant human *in vitro* platform for studying CNS infection (*33*).

Herein, we established a human cerebral organoid model of WNV_NSW2011_ infection and used it to elucidate the cellular basis of WNV neuropathogenesis. Using virus-inclusive single-cell RNA sequencing (scRNA-Seq) of the infected organoids, we mapped WNV tropism across neural cell types, characterised cell-type-specific innate immune responses and identified host determinants of differential susceptibility. We show that WNV preferentially infects neural progenitors, astrocytes, and early neurons, while mature neurons exhibited a profound resistance to infection. This tropism was confirmed by immunostaining of infected organoids and virus replication assays in monocultures of non-malignant human neural cells. In infected cerebral organoids, astrocytes were found to act as the principal innate immune responders. Progenitor cells also demonstrated strong activation of antiviral defense pathways, while neurons showed limited reaction to infection and interferon (IFN). This was found to be consistent with low expression of pattern-recognition receptors and interferon signalling components in mature neurons. Given that cell-specific differences in innate immune responses cannot explain viral tropism to immune-responsive neural progenitors and glial cells, we hypothesised that WNV-susceptible cells may express host dependency factor that is lacking in WNV-resistant mature neurons. Using the differential gene expression test between infected and uninfected cells within the organoid tissue we identified the serine biosynthesis enzyme phosphoglycerate dehydrogenase (PHGDH) as a putative host dependency factor for WNV neuro-infection. We confirmed the requirement of PHGDH for WNV replication using RNAi knock-down and pharmacological inhibition of PHGDH. Finally, we demonstrated the antiviral activity against WNV of the FDA-approved PHGDH-antagonising drug disulfiram in cultured human neural progenitors and brain organoids. These findings provide a human-relevant model of WNV neuropathogenesis, reveal specific functions of different neural cell populations in antiviral protection of brain tissue and identify PHGDH as a potential therapeutic target for host-directed treatment of WNV encephalitis.

## RESULTS

### WNV_NSW2011_ productively infects human cerebral organoids

To establish the organoid model of WNV infection we generated cerebral organoids from the human iPSC line WTC-11 using a modified suspension-based protocol (*34*). The organoids were inoculated at two maturation stages, 40 days *in vitro* (DIV40) and 60 days *in vitro* (DIV60), with 10^4^ or 10^5^ focus-forming units (FFU) of WNV_NSW2011_ per organoid. Viral titers were then determined in culture fluid samples every 2 days over a 21-day time course. WNV_NSW2011_ replicated productively in organoids of both ages and at both inoculation doses, with titers rising rapidly within the first two days of infection and reaching plateau at approximately 10^5^ FFU/ml for the remainder of the experiment (Fig. 1, A, B). The two infection doses produced comparable titers that converged within the first 3 days. However, the replication kinetics of individual organoids were more uniform at the higher inoculation dose (Fig. 1, A, B, right panels). At 10^5^ FFU, organoid-to-organoid variation was minimal throughout the entire time course, whereas the 10^4^ FFU dose yielded greater variability in the early (1-5 days) and late (12-21 days) phases of infection (Fig. 1, A, B, right panels). Therefore, we selected 10^5^ FFU as the standard inoculation dose for all subsequent experiments.

**Fig. 1.**
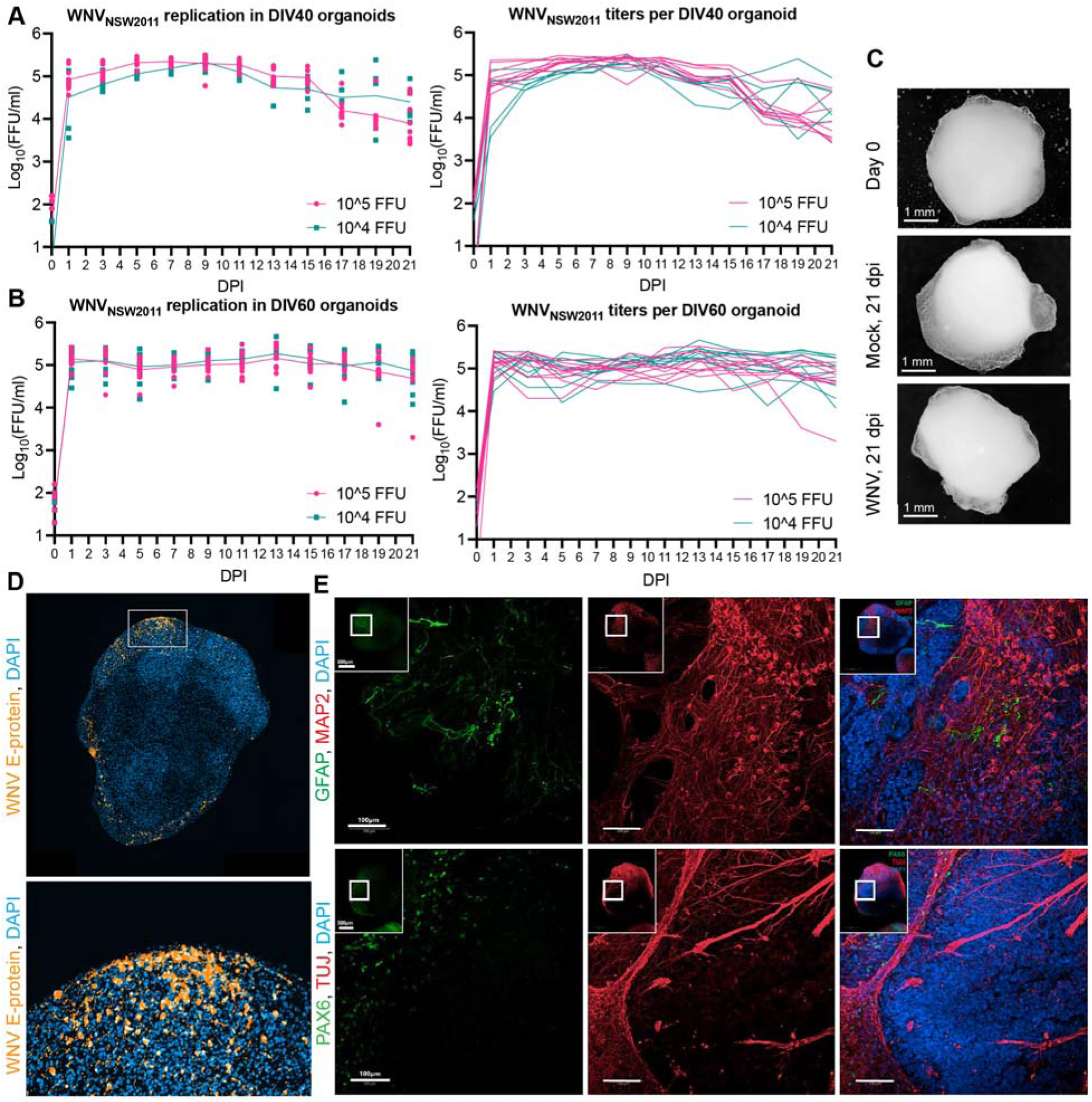
WNV_NSW2011_ productively infects human cerebral organoids. **(A, B)** Replication of WNV_NSW2011_ in human cerebral organoids at **(A)** 40 days in vitro (DIV40) and **(B)** 60 days in vitro (DIV60). Organoids were inoculated with 10 or 10 FFU of WNV_NSW2011_ and culture supernatants were collected every 2 days through 21-day time course. Viral titers were determined by immunofluorescent focus-forming assay on C6/36 cells. Left panels show mean viral titers per group with individual data points; right panels show individual virus replication kinetics for each organoid. DIV40: n=9 (10 FFU) and n=6 (10 FFU); DIV60: n=10 per group. Organoids were derived from 2 independent differentiation batches of the WTC-11 iPSC line. **(C)** Representative bright-field images of DIV60 cerebral organoids before infection (Day 0) and of mock- and WNV_NSW2011_-infected organoids at 21 dpi after inoculation with 10^5^ FFU of WNV_NSW2011_. Scale bars are 1 mm. **(D)** Immunofluorescent detection of WNV envelope (E) protein (orange) in sections of DIV60 cerebral organoids inoculated with 10 FFU of WNV_NSW2011_ at 3 dpi. Sections are counterstained with DAPI (blue). The boxed region is shown at higher magnification below. **(E)** Immunofluorescent staining of sections from uninfected DIV60 cerebral organoids for the markers of astrocytes (GFAP), radial glia (PAX6), immature neurons (TUJ1) and post-mitotic neuron marker (MAP2); nuclei counterstained with DAPI. Boxed regions are shown as higher-magnification images. Scale bars are 100 μm. Images in (C-E) are representative of at least three organoids that showed similar results.

Bright-field imaging of the infected organoids showed that they remained morphologically intact throughout the 21-day time course without signs of disintegration (Fig. 1C) unlike what we previously observed with ZIKV-infected organoids (*29, 30*). *In situ* infection in organoid tissue was confirmed by immunohistochemical detection of WNV envelope (E) protein at 3 days post-infection (dpi), which revealed areas of infected cells primarily in the outer layer of the organoid parenchyma (Fig. 1D).

We further selected DIV60 organoids as the primary model for downstream analyses given that at this more advanced stage of maturation the organoids are expected to harbor a broader diversity of neural cell types. The immunofluorescent staining of uninfected organoids for the markers of neural cell types revealed the presence of PAX6-positive radial glia (progenitor cells), GFAP-positive astrocytes, TUJ1-positive immature neurons, and MAP2-positive post-mitotic neurons (Fig. 1E), demonstrating that the model contains the major neural cell populations relevant to WNV neuropathogenesis. Together, these data demonstrate that DIV60 human cerebral organoids have cellular heterogeneity and consistently sustain WNV_NSW2011_ replication.

### WNV preferentially infects neural progenitors, early neurons and astrocytes, while mature neurons are largely resistant to infection

To identify the cellular and molecular determinants of WNV neural infection, we performed single-cell transcriptomic profiling of infected cerebral organoids. A major technical challenge for single-cell analysis of flavivirus infection is that the viral genome lacks a 3’ poly(A) tail. Widely used droplet-based scRNA-Seq platforms, including standard 10x Genomics 3’ Chromium workflow, rely on oligo(dT)-mediated capture of polyadenylated host mRNAs and therefore fail to reliably detect non-polyadenylated viral RNA (*35*). In addition, these protocols require the use of live unfixed cells (*36*), which poses a biosafety risk when working with the infected material. To overcome these limitations, we utilised a 10x Genomics Chromium Next GEM Single Cell Fixed RNA protocol, which doesn’t involve poly(A)-capture and relies on gene-specific probes for library preparation from formaldehyde-fixed cells (*37*). To enable simultaneous capture of viral and host transcripts in the scRNA-Seq experiment we designed three custom probe pairs that bind to the WNV genomic RNA (Fig. 2A, Supplementary table S1). The WNV-specific probes were added as spike-in into pan-transcriptomic probe cocktail provided with the 10x Genomics reagents kit during scRNA-Seq library preparation.

**Fig. 2.**
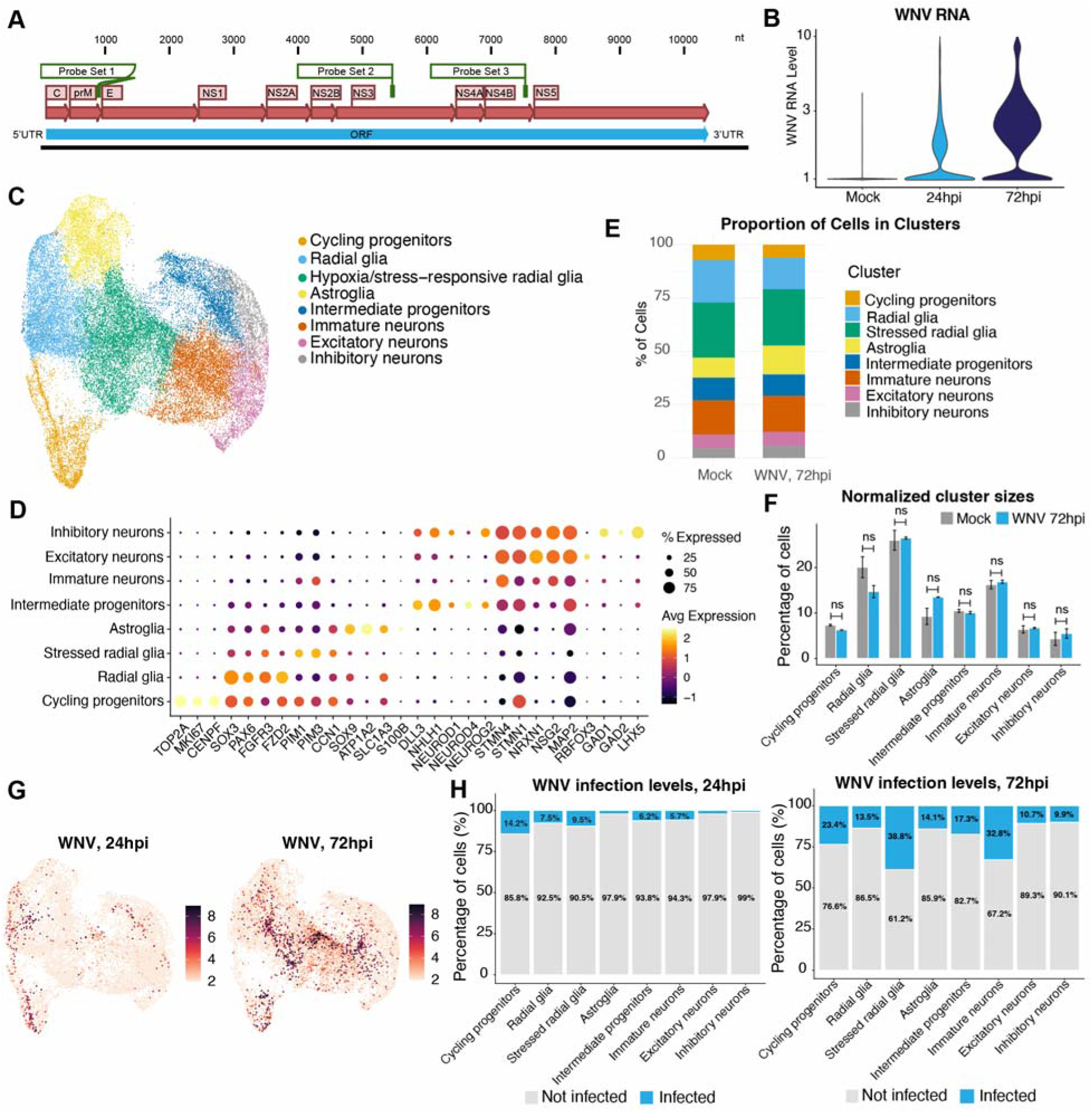
Virus-inclusive scRNA-Seq of WNV-infected cerebral organoids demonstrates preferential viral tropism to immature and glial cells. (**A**) Location of custom probe binding sites across the WNV genome used for virus-inclusive single cell gene expression profiling. (**B**) Per-cell WNV RNA abundance determined by virus-inclusive scRNA-seq, in mock- and WNV_NSW2011_-infected organoids at 24 and 72 hpi. Groups of DIV60 organoids were inoculated overnight with 10 FFU of WNV_NSW2011_. At the indicated time points organoids were pooled (eight per pool), dissociated to single-cell suspensions, and ∼10,000 cells per sample were processed with the 10x Genomics Chromium Next GEM Single Cell Fixed RNA kit using a custom probe spike-in workflow. Data are from two biological replicates. (C) Uniform manifold approximation and projection (UMAP) visualisation of the single-cell gene expression profile of human cerebral organoids. Each point represents one cell, coloured according to annotated cell type. Cell clusters were identified by unsupervised graph-based clustering and annotated based on marker gene expression. **(D)** Expression of marker genes for cycling neural progenitors (*TOP2A*, *MKI67*, *CENPF*), radial glia (*SOX3*, *PAX6*, *FGFR3*, *FZD2*), astrocytes (*SOX9*, *ATP1A2*, *SLC1A3*, *S100B*), intermediate progenitors (*DLL3*, *NHLH1*, *NEUROD1*, *NEUROD4*, *NEUROG2*), post-mitotic neurons (*STMN4*, *STMN1*, *NRXN1*, *NSG2*, *MAP2*, *RBFOX3*), and inhibitory neurons (*GAD1*, *GAD2*, *LHX5*). *PIM1*, *PIM3*, and *CCN1* indicate stress- and hypoxia-associated transcriptional states. Visualisations in (C) and (D) are based on cumulative, non-integrated data from mock- and WNV-infected organoids across two replicates. **(E)** Relative proportions of each annotated cell type in mock-versus WNV-infected (72 hpi) organoids (pooled data). **(F)** Per-cluster cell-type proportions in mock-versus WNV-infected (72 hpi) organoids; bars are the mean of two independent experiments ± SEM. **(G)** Feature plots showing detection of WNV RNA in single cells from infected human cerebral organoids at 24 and 72hpi. Cells are displayed on the UMAP embedding generated from single-cell transcriptomes, with colour intensity representing the abundance of WNV RNA detected in each cell. **(H)** Proportion of WNV RNA-positive cells in each annotated cell cluster at 24 and 72hpi, defined as cells containing at least two unique molecular identifiers (UMIs) from WNV-derived reads. Data in (G) and (H) are cumulative from two independent experiments.

To perform single-cell gene expression analysis, groups of eight DIV60 cerebral organoids were inoculated in duplicates with 10^5^ FFU of WNV_NSW2011_ and collected at 24 and 72hpi. Mock-infected organoids were used as a control. Infection of the organoids was confirmed by virus titration (Fig. S1A). At the indicated time points the organoids from each group were pooled and dissociated into single-cell suspensions. Cells were fixed with 4% formaldehyde and analysed by virus-inclusive scRNA-Seq. The scRNA-Seq detected accumulation of WNV RNA in cells derived from infected organoids at 24 and 72 hpi, with minimal signal in mock-infected organoids, confirming specific and sensitive detection of viral RNA (Fig. 2B). The resulted single-cell transcriptomic profiles showed high consistency between the replicates (Fig. S1B), whereas the differences in UMAP-projections were observed between infected and uninfected group, indicating for the transcriptional changes induced by infection (Fig. S1C). Unsupervised clustering of single-cell transcriptomes resolved eight distinct cell populations (Fig. 2C) that were all present in each of the analysed groups (Fig. S1D). Cell identities in the clusters were annotated by canonical marker gene expression (Fig. 2C, D, Fig. S1E, F). The organoids contained neural progenitor cells including cycling progenitors (*TOP2A*^+^, *MKI67*^+^, *CENPF*^+^) and radial glia (*SOX3*^+^, *PAX6*^+^, *FGFR3*^+^, *FZD2*^+^) (Fig 2C, D, Fig S1F). Of the neuronal differentiation axis (*STMN4*^+^), the organoids contained intermediate progenitors (*DLL3*^+^, *NHLH1*^+^, *NEUROD1*^+^, *NEUROD4*^+^, *NEUROG2*^+^) and post-mitotic neurons (*MAP2*^+^, *NSG2*^+^), including mature neurons (*RBFOX3*^+^, high *STMN1*, *NRXN1,* mature synapses *CPLX1^+^*) with a distinct population of inhibitory neurons (*GAD1*^+^, *GAD2*^+^, *LHX5*^+^) as well as immature neurons negative for *RBFOX3* (NeuN) and expressing low levels of *NSG2*, *NRXN1* and *STMN1* (Fig. 2C, D, Fig. S1F). Within the astroglial cluster, ∼20-25% of cells were mature astrocytes (*S100B*^+^, *SPARCL1*^+^), while the remainder were still maturing (Fig. 2C, D, Fig. S1F) although clearly committed to the astroglial lineage (*SOX9*^+^, *ATP1A2*^+^, *SLC1A3*^+^). A separate cluster contained cells co-expressing radial glia markers and hypoxia-response/stress genes (e.g., *PIM1*, *PIM3*), likely corresponding to radial glia in the organoid core where limited oxygen diffusion induces hypoxic stress (Fig. 2C, D). The relative proportions of these populations were largely comparable between mock- and WNV-infected organoids (Fig. 2E, F). At 72hpi we detected slight increase in size of the astroglia cluster, which may indicate for proliferation of activated astrocytes (Fig. 2E). However, this difference was not statistically significant in comparison to mock (Fig. 2F). This indicates that, unlike Zika virus (ZIKV) (*29*), WNV doesn’t cause the depletion of any specific cell types, at least at early infection time points. The observation is consistent with the lack of organoid disintegration upon WNV (Fig. 1C), but not ZIKV (*29, 30*) infection and points to the potential differences in neuropathogenesis of these viruses.

To characterize cell tropism of WNV we explored presence of viral genomic RNA in different cell clusters. To exclude potential ambient RNA contamination and cells carrying viral RNA without productive infection (e.g., surface-attached virus), we defined cells as WNV-infected only if they contained at least two WNV-derived unique molecular identifiers (UMIs), as was previously applied for scRNA-seq analysis of mouse brain infected with Japanese encephalitis virus (JEV) (*38*). At 24 hpi very few infected cells were detected (Fig. 2G). They were located primarily in the clusters of cycling progenitors, radial glia, intermediate progenitors and immature neurons (Fig. 2G, H). Mature neurons and astrocytes were largely uninfected at this time point (Fig. 2G, H). At 72hpi, the number of infected cells increased. Approximately 39% of stressed radial glia cells contained moderate to high levels of viral RNA, comprising the most infected cluster (Fig. 2G, H). Other progenitor cell clusters were also highly infected (Fig. 2G, H). At this time point viral RNA was also detected in astroglia (14.1%) (Fig. 2G, H). Mature neuron clusters (excitatory and inhibitory neurons) had the lowest presence of infected cells (∼10%) (Fig. 2H) and also contained low levels of viral RNA (Fig. 2G). These results suggest that in human cerebral organoids WNV has preferential tropism to immature and glial cells and low tropism to mature neurons.

The viral tropism identified by scRNA-Seq was further corroborated by immunofluorescent staining of infected organoid sections, in which WNV E protein localized extensively in the cells positive for the neural progenitor marker SOX2 and the immature neuron marker DCX (Fig. 3A). Small number of cells expressing the astrocyte marker S100B were also positive for WNV (Fig. 3A). All cells expressing the mature neuron marker NeuN were uninfected (Fig. 3A). To test whether this susceptibility pattern reflects the intrinsic characteristics of neural cell types rather than a property of the organoid environment, we infected 2D cultures of human neural cells. The panel contained immortalised non-malignant human astrocytes SVG p12, neural progenitor cells ReNcell VM and post-mitotic human neurons that was obtained by differentiation of ReNcell VM (Fig. S2). We also assessed viral replication in immortalised human microglial cells HMC3 to extend our analysis to a CNS-resident cell type that is absent from standard cerebral organoids due to its non-neuroectodermal origin. Virus growth kinetics analysis demonstrated that WNV_NSW2011_ replicated efficiently in neural progenitors and astrocytes, while in ReNcell-derived mature neurons the viral titers remained at the level of the residual inoculum observed at 0 hpi (∼10^2^ FFU/ml) (Fig. 3B). WNV_NSW2011_ also did not replicate to a meaningful level in HMC3 microglia cells (Fig. 3B). The same pattern was observed for the prototypic North American strain WNV_NY99_ (Fig. 3C), indicating that preferential infection of progenitor and astroglial cells over mature neurons is a conserved property of WNV and not a specific characteristic of the NSW2011 strain. Collectively, these results identify neural progenitors, astrocytes and immature neurons as the principal cellular targets of WNV in the human neural tissue, with mature neurons being largely resistant to infection.

**Fig. 3.**
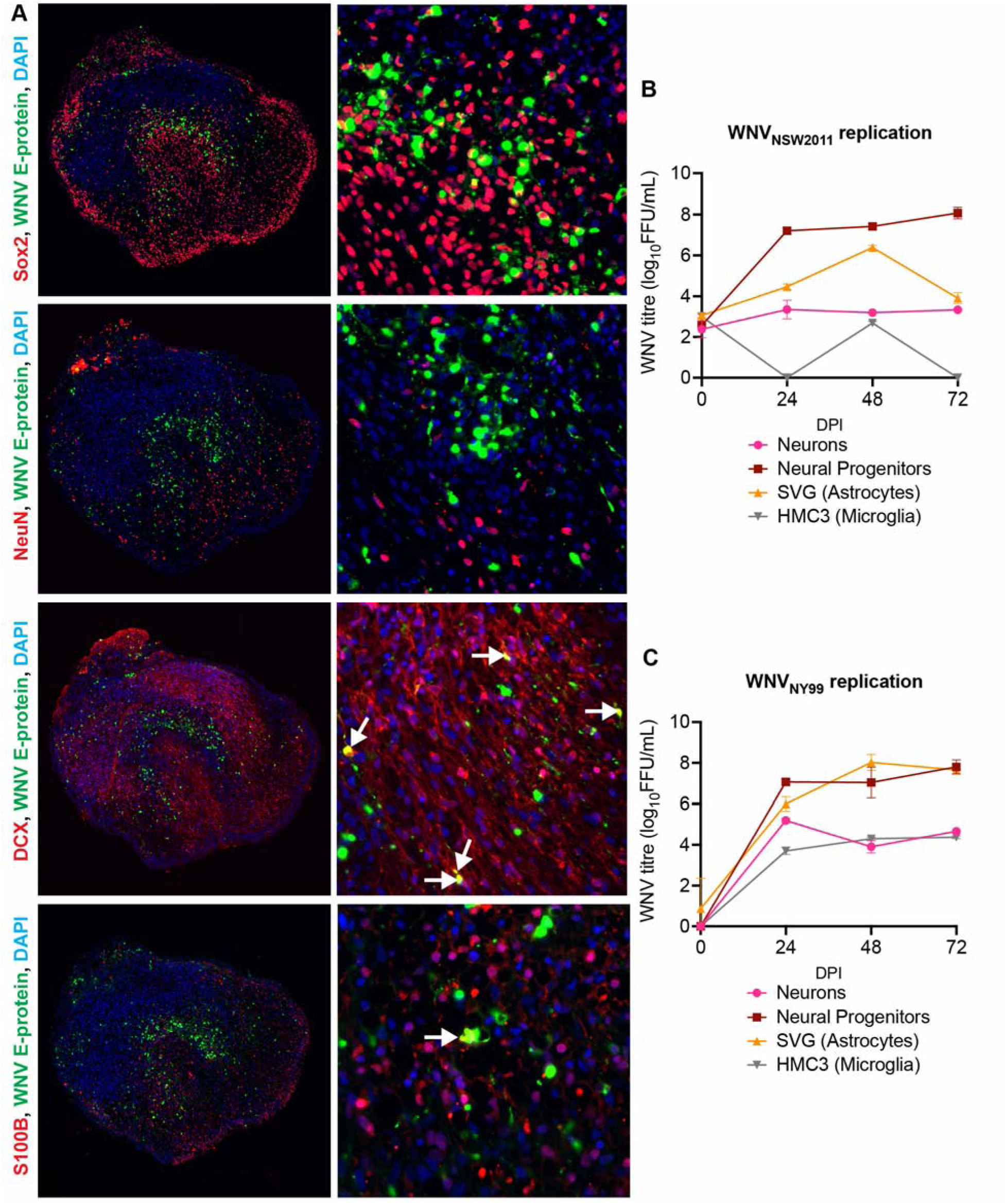
Immunostaining of infected cerebral organoids and virus replication analysis in neural cell culture models indicate preferential WNV infection of progenitor and astroglial cells over mature human neurons. (**A**) Immunofluorescent staining of the histological sections of WNV_NSW2011_-infected human cerebral organoid showing detection of WNV envelope (E) protein together with markers of neural progenitors (SOX2), mature neurons (NeuN), immature neurons (DCX), and astrocytes (S100B). Left panels show low-magnification images of whole organoid sections; right panels show higher-magnification views of infected regions. Arrows indicate WNV E-positive cells co-localising with the indicated cell type markers. **(B)** Replication of WNV_NSW2011_ in cultures of human neural cells. **(C)** Replication of WNV_NY99_ in cultures of human neural cells. Cells in **(B, C)** were infected with WNV at MOI=1 and viral titers in culture fluids were determined by focus-forming assay on C6/36 cells. Enriched neuronal populations (Neurons in **B** and **C**) were produced by differentiation of ReNcell VM neural progenitor cell line through growth factor withdrawal and infected at day 12 of differentiation. Data are presented as means from three biological replicates ± SD.

### Astrocytes are the principal innate immune responders to WNV infection

We hypothesised that the differential cell tropism of WNV in neural tissue may be determined by the differences in antiviral responses between cell populations. Therefore, we next examined the innate immune response to WNV infection across organoid cell types. Computational intercellular communication inference based on single cell gene expression analysis predicted astroglia to be the key hub of type I interferon signalling in the infected organoids, acting as a primary sender and as a receiver of signal (Fig. 4A). Closer examination of IFN-expressing cells revealed only marginal expression of IFNs at 24 hpi (Fig. 4B; S3A). At 72 hpi the number of IFN-expressing cells had increased yet remained very low given the abundance of the infected cells at this time point (Fig. 1G, H). The small number of cells responding to infection by IFN production is consistent with prior findings in cultured HEK293 cells (*39*) and indicates that WNV efficiently evades sensing by pattern recognition receptors (PRRs) or inhibits downstream signalling required for activation of the IFN genes. The type I interferon *IFNB1* and the type III interferon *IFNL1* were the most highly expressed interferon genes, as evident from transcript levels and the number of expressing cells (Fig. 4B; S3A, B). At this time point, the IFN-producing cells were concentrated in the astroglia cluster (Fig. 4B) with few scattered *IFNB1*-expressing cells also present among stressed radial glia, intermediate progenitors and immature neurons (Fig. 4B). These results identify astrocytes as main sensors of WNV infection in human cerebral organoids that drive antiviral response in the tissue through production of IFNB1 and IFNL1.

**Figure 4.**
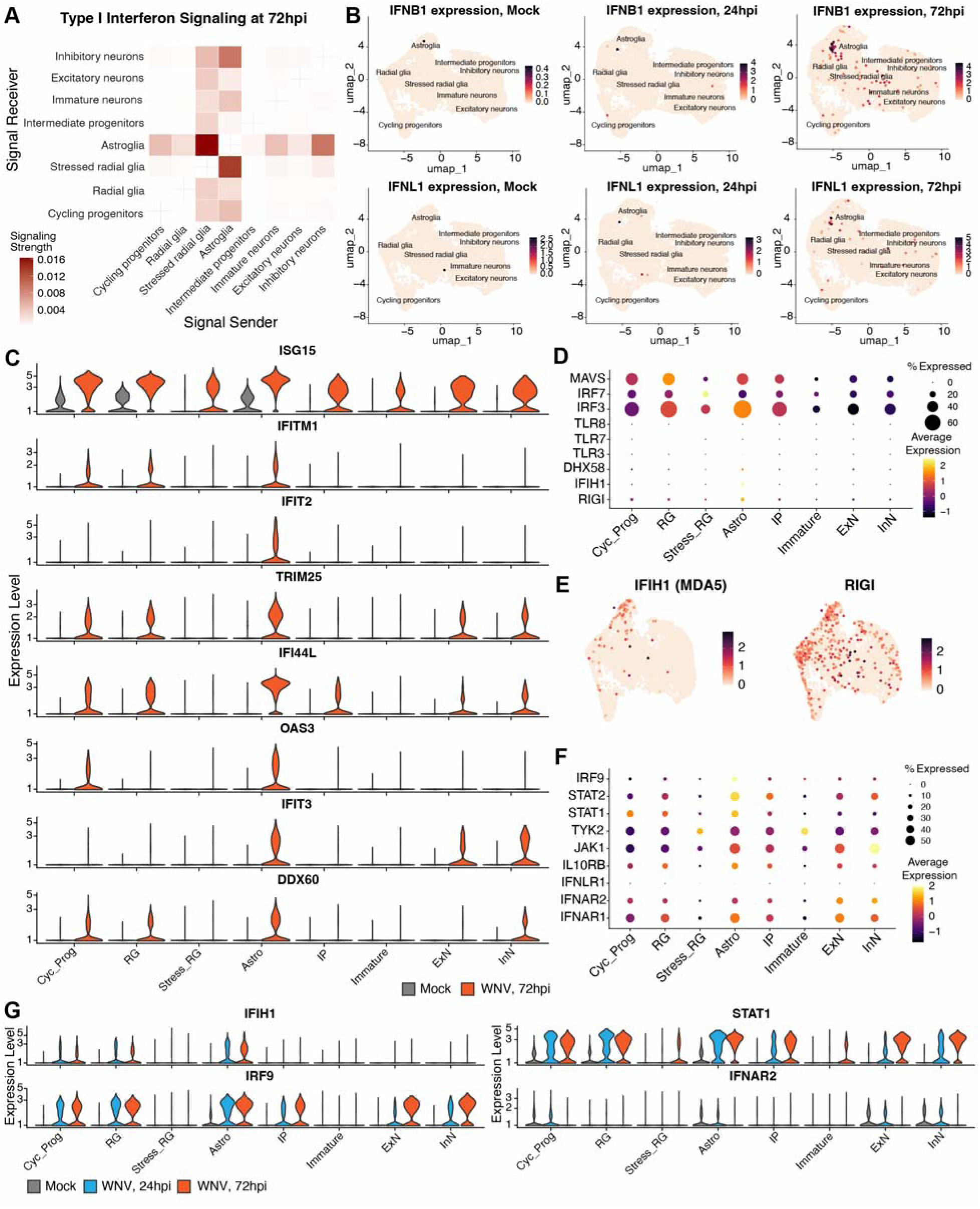
Innate immune responses to WNV infection in human cerebral organoids are cell-type-specific. **(A)** Inferred type I interferon signalling between cell populations in WNV-infected cerebral organoids at 72 hours post-infection (hpi). Rows indicate predicted signal-receiving cell populations, columns indicate predicted signal-sending populations, colour intensity represents signalling strength. **(B)** Expression of *IFNB1* and *IFNL1* in mock-infected and WNV-infected cerebral organoids at 24 and 72 hpi. **(C)** Expression of representative interferon-stimulated genes (ISGs) across the cell populations in mock-infected and WNV-infected cerebral organoids at 72 hpi. **(D)** Aggregated expression of viral RNA sensors and signalling components of the PRR-pathways across cell populations in WNV-infected organoids at 72 hpi. Dot size indicates the percentage of cells expressing each gene, and colour indicates average normalised expression. **(E)** UMAP feature plots showing expression of the viral RNA sensors MDA5 (*IFIH1*) and RIG-I (*DDX58*) in individual cells of uninfected cerebral organoids. **(F)** Expression of interferon receptors and downstream interferon signalling genes across cell populations in WNV-infected organoids at 72 hpi. Dot size indicates the percentage of cells expressing each gene, and colour indicates average normalized expression. **(G)** Aggregated expression of interferon receptors and signalling genes across cell populations in mock-infected and WNV-infected organoids at 24 and 72 hpi. Cells in (B, E) are displayed on the UMAP embeddings, each point represents one cell, colour intensity indicates normalised gene expression levels. Cyc_Prog - cycling progenitors, RG - radial glia, Stress_RG - stressed radial glia, Astro - astroglia, IP - intermediate progenitors, Immature - immature neurons, ExN - excitatory neurons, InN - inhibitory neurons.

Despite the small number of IFN-expressing cells, secreted interferons elicited robust upregulation of interferon-stimulated genes (ISGs), including *ISG15*, *IFITM1*, *IFIT2*, *TRIM25*, *IFI44L*, *OAS3*, *IFIT3*, and *DDX60* (Fig. 4C; S3C, S4). However, the ISG response was uneven across cell types. The fastest and highest upregulation of ISGs was observed in astroglia and progenitor populations, while neurons exhibited delayed and week response to IFN (Fig. 4C; S3C). Among the selected ISGs, only *ISG15* showed moderate induction in neurons at 24 hpi, whereas astrocytes displayed strong up-regulation of all antiviral genes in the panel (Fig. S3C). Progenitor cells also showed up-regulation of multiple ISGs at this time point (Fig. S3C). At 72 hpi, expression of *TRIM25*, *IFI44L* and *IFIT3* was observed in mature neurons, although at lower level than in astrocytes (Fig. 4C). The expression of *IFITM1*, *IFIT2* and *OAS3* was still lacking in neurons at this time point, while *DDX60* was upregulated in small number of inhibitory neurons (Fig. 4C). In the immature neurons, the antiviral response was also limited to *ISG15* up-regulation (Fig. 4C). The transcriptome-wide pseudo-bulk gene expression comparison of individual cells clusters between infected and uninfected organoids also identified astrocytes and radial glia as the cell populations having the strongest response to WNV as indicated by the number of differentially expressed genes (DEGs) and magnitude of change in their expression, while only few DEGs were identified in mature neurons (Fig. S4). In addition, the Gene Ontology and pathway enrichment analyses of the astroglial cluster revealed strong transcriptional signatures associated with antiviral response in these cells in the infected organoids at 24 and 72 hpi (Fig. S5). Together, these data indicate that astrocytes mount the principal innate immune response to WNV in human cerebral organoids alongside the progenitor cells that also develop a strong IFN response, while mature neurons respond weakly and late.

### Neurons lack expression of pattern-recognition receptors and interferon signalling component STAT1

To obtain a mechanistic explanation of the observed cell-specific differences in IFN production and response, we examined the expression of innate immune sensors and IFN signalling components across organoid cell types. The viral RNA-sensing machinery includes the pattern-recognition receptors (PRRs) RIG-I (*DDX58*), MDA5 (*IFIH1*), and LGP2 (*DHX58*), the endosomal Toll-like receptors *TLR3*, *TLR7*, and *TLR8*, the signalling adaptor MAVS, and the transcription factors IRF3 and IRF7 (*40*). For detection of WNV specifically, the cooperative action of RIG-I and MDA-5 is known to be required (*41, 42*). In uninfected cerebral organoids MDA5 was expressed almost exclusively in astrocytes, while RIG-I expression was detected in astrocytes and progenitor cells (Fig. 4D, E). Neurons also had low expression of other components of the PRR-signalling pathway (Fig. 4D). At 24 and 72 hpi, expression of RIG-I and MDA5 increased and was present in small number of cells within the neuronal clusters (Fig. S3D). However, unlike in astrocytes and progenitor cells, infection did not lead to the substantial overall boost in their expression across the cluster (e.g. *IFIH1* in Fig. 4G). Thus, the limited IFN response in neurons can be attributed to low baseline expression of viral RNA sensors and signalling components, which, unlike in astrocytes and progenitors, are not robustly induced upon infection.

The downstream type I and III interferon signalling cascades consist of the interferon receptors (IFNAR1, IFNAR2, IFNLR1) and the downstream Janus kinase/signal transducer and activator of transcription (JAK-STAT) signalling components JAK1, TYK2, STAT1, STAT2, and IRF9 (*43*). Of these signalling components, the type III IFN receptor IFNLR1 was not expressed in any cell population in uninfected organoids and was not induced upon infection (Fig. 4F; S3E). This indicates that type III IFN is unlikely to contribute to the antiviral response in this system. The two subunits of the type I IFN receptor, IFNAR1 and IFNAR2, were expressed in most cell populations (Fig. 4F; S3E), suggesting that IFNB1 (Fig. 4B) likely serves as the principal mediator of defence against WNV in neural cells.

Among the downstream signalling components, JAK1, TYK2, and STAT2 were expressed ubiquitously across all cell types in uninfected organoids, mostly at low-to-moderate levels Fig. 4F). However, STAT1 expression was restricted to cycling progenitors, radial glia, and astrocytes (Fig. 4F). Astroglia showed the highest basal expression of IFN signalling genes overall (Fig. 4F). Following WNV infection, all components of the IFN pathway were strongly up-regulated in most cell types apart from stressed radial glia and immature neurons (Fig. 4G; S3E). Consistent with the delayed response of the mature neurons to IFN (Fig. 4C, S3C), they only reached high STAT1 levels by 72 hpi (Fig. 4G). By this time, however, IFNAR2 had been downregulated (Fig. 4G), consistent with engagement of the known negative feedback loop that limits innate immune overactivation (*44*). The late induction of STAT1 in neurons therefore coincides with loss of receptor availability, which likely explains their delayed and weak IFN response.

Overall, the analysis of innate immune responses across organoid cell types identified neurons and stressed radial glia as having low antiviral response competency, which is consistent with their lack of basal expression and induction of RNA sensing and IFN signalling components. This deficiency can explain permissiveness to WNV infection of stressed radial glia and immature neurons (Fig 2G, H). However, mature neurons that are also deficient in viral sensing and have a limited capacity to respond to interferons are resistant to infection, unlike the IFN-competent astrocytes and progenitor cells. Therefore, innate immune competency cannot fully explain the observed cell-specific tropism of WNV to immature and glial cells vs mature neurons (Fig 2G, H).

### PHGDH is a host dependency factor for WNV infection of susceptible neural cells

Given that the differences in innate immune responses cannot fully explain WNV tropism in human cerebral organoids, we hypothesised that cell-specific host factors may determine which cells support viral replication. To identify such factors, we performed differential gene expression analysis between highly WNV-positive (>4UMI) and WNV-negative cells within infected organoids. The analysis identified a set of genes significantly enriched in infected cells (FDR-adjusted *p*-value < 0.05). Of them, the most strongly associated with WNV-infected cells were the genes encoding phosphoglycerate dehydrogenase (*PHGDH*), eukaryotic translation initiation factor 4E-binding protein 1 (*EIF4EBP1*) and tribbles pseudokinase 3 (*TRIB3*), together with serine hydroxymethyltransferase 2 (*SHMT2*) and methylenetetrahydrofolate dehydrogenase 2 (*MTHFD2*) (Fig. 5A). Among the three most significantly associated genes, *EIF4EBP1* and *TRIB3* (both are ATF4-regulated stress-responsive genes) were induced upon infection, whereas *PHGDH* levels remained largely unchanged between mock- and WNV-infected cells across the clusters (Fig. 5B). The infection-independent expression pattern of *PHGDH* together with its specific association with WNV-permissive cells suggest that it may represent an intrinsic cellular feature predisposing cells to WNV infection, rather than a downstream consequence of the infection or antiviral responses.

**Fig. 5.**
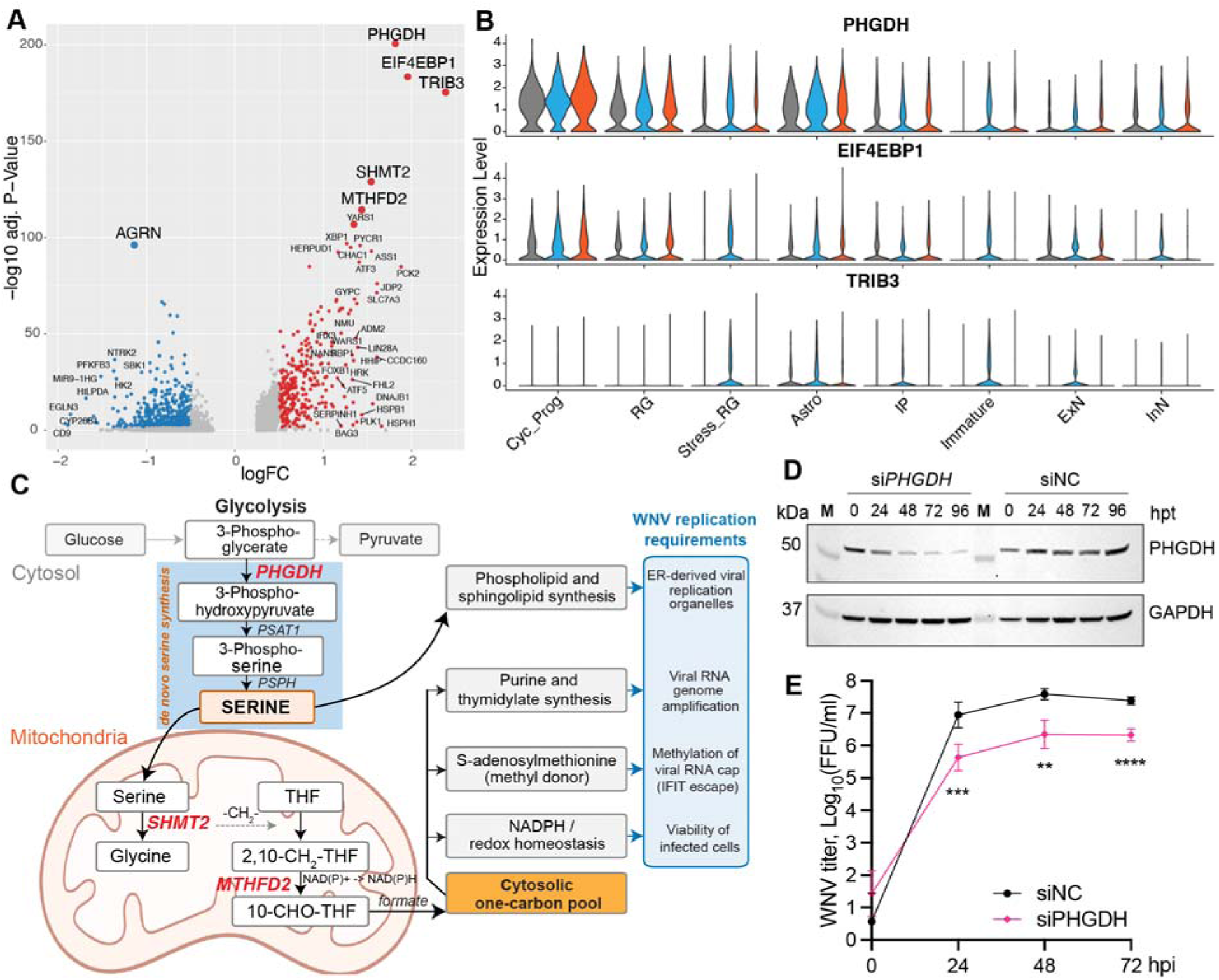
Single-cell transcriptomic analysis identifies PHGDH as a host factor for WNV replication in susceptible neural cells. **(A)** Genes expression signatures associated with WNV-positive cells in infected human cerebral organoids. Genes significantly (FDR-adjusted p-value <0.05) enriched in WNV RNA-positive cells are shown in red, and genes depleted in WNV RNA-positive cells are shown in blue. **(B)** The expression of the top differentially expressed genes associated with WNV-positive is not induced in response to infection. **(C)** Schematic of *de novo* serine synthesis and mitochondrial one-carbon metabolism. PHGDH catalyses the committed, rate-limiting diversion of the glycolytic intermediate 3-phosphoglycerate into serine biosynthesis. Mitochondrial SHMT2 transfers the serine-derived one-carbon unit to tetrahydrofolate (THF), and MTHFD2 oxidises 5,10-methylene-THF to 10-formyl-THF, sustaining the cytosolic one-carbon pool. Outputs of the pathway (purine and thymidylate synthesis, S-adenosylmethionine-dependent methylation, NADPH-dependent redox balance) and serine-derived phospholipid/sphingolipid synthesis are each required for flavivirus replication. Enzymes whose transcripts are enriched in WNV RNA-positive cells are highlighted in red. **(D)** Western blotting detection of PHGDH expression in ReNcell VM cells transfected with siRNAs. siNC – negative control siRNA. GAPDH is shown as a loading control. **(E)** Replication of WNV_NSW2011_ in cells transfected with *PHGDH* siRNA. Cells were infected at MOI=0.1 at 24 hpt. Viral titers were determined by immunofluorescent focus-forming assay on C6/36 cells. Statistical analysis is by one-way ANOVA with Sidak’s correction; n=3, **p=0.0018, ***p=0.0008, ****p<0.0001. The values are the means ± SD.

PHGDH catalyses the rate-limiting step of de novo serine biosynthesis, generating serine that feeds into the one-carbon metabolism cycle to supply precursors for nucleotide synthesis, methylation, and redox homeostasis (Fig. 5C). Notably, two other genes associated with WNV-infected cells, *SHMT2* and *MTHFD2*, encode downstream enzymes of the same pathway (Fig. 5C). The simultaneous high expression of multiple one-carbon pathway enzymes in WNV-permissive cells suggests that baseline flux through serine/one-carbon metabolism may represent a condition required for WNV susceptibility and identifies PHGDH, SHMT2 and MTHFD2 as potential host dependency factors for WNV.

To test the requirement of *PHGDH* expression for WNV replication in human neural cells, we assessed WNV replication in neural progenitor cells following RNAi-mediated PHGDH depletion (Fig. 5D, E). The siRNA-mediated knockdown of PHGDH (Fig. 5D) significantly reduced WNV_NSW2011_ replication compared to the cells transfected with a non-targeting control siRNA (Fig. 5E). In addition, treatment of human neural progenitors with the small-molecule PHGDH inhibitor CBR-5884 reduced WNV_NSW2011_ replication by up to 2.5-log10 relative to vehicle-treated control (Fig. 6A), providing further evidence for its requirement for viral replication. Lack of the cytotoxicity of the compound in neural progenitor cells (Fig. 6B) indicates that the observed effect is due to PHGDH suppression rather than drug-induced cell death. Together with the results of scRNA-Seq, these data identify PHGDH, and *de novo* serine biosynthesis more broadly, as a host dependency pathway for WNV in susceptible neural cells and nominate it as a candidate target for host-directed antiviral intervention.

**Fig. 6.**
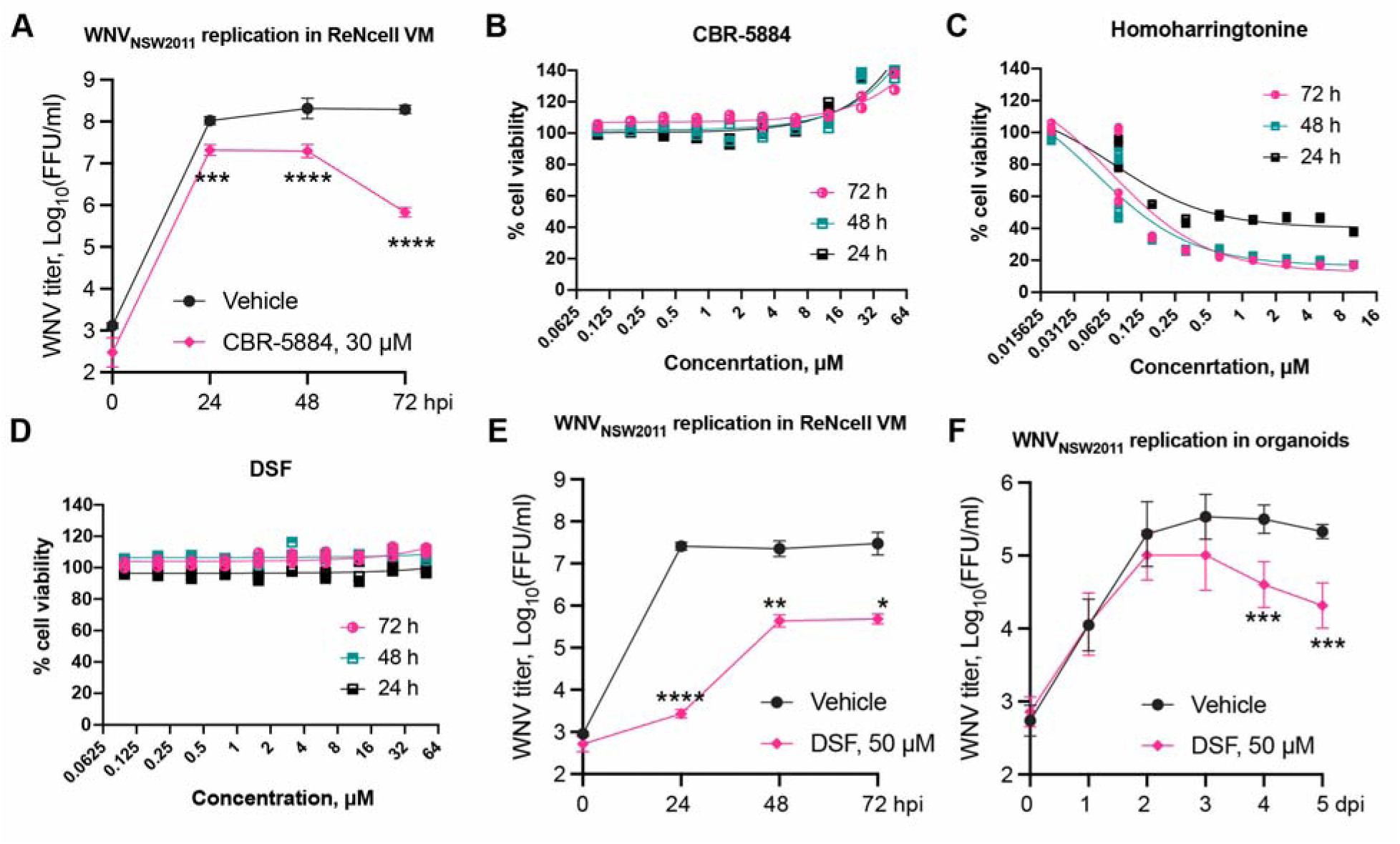
Pharmacological inhibition of PHGDH reduces WNV replication. **(A)** Replication of WNV_NSW2011_ in ReNcell VM cells treated with CBR-5884 small molecule PHGDH inhibitor. **(C-D)** The effect of the drugs used in the experiment on viability of ReNcell VM cells. Cell viability was assessed after treatment with CBR-5884 **(B)**, Hommoharringtonine **(C)** and Disulfiram **(D)** using CellTiter 96 AQueous One Assay (Promega) and expressed as percentage compared to untreated control. **(E)** Replication of WNV_NSW2011_ in ReNcell VM cells treated with Disulfiram (DSF). (F) Replication of WNV_NSW2011_ in human cerebral organoids cells treated with Disulfiram (DSF). Cells in **(A)** and **(E)** were infected at MOI=0.1. Cerebral organoids in **(F)** were inoculated with 10^5^ FFU of WNV. In **(A-E)** n=3 and in **(F)** n=5. Statistical analysis in **(A, E, F)** is by repeated measures one-way ANOVA with Sidak’s correction; *p<0.05, **p<0.01, p***p<0.001, ****p<0.0001.

### The FDA-approved PHGDH-inhibitor disulfiram reduces WNV replication in human neural progenitor cultures and cerebral organoids

Having established that the PHGDH inhibitor CBR-5884 reduces WNV replication, we next explored whether pharmacological agents already approved for human use could also target PHGDH. Such activity has been reported for homoharringtonine (*45*), a plant alkaloid approved for cancer treatment, and for disulfiram (DSF) (*46*), used in the management of alcohol abuse disorder through its inhibition of aldehyde dehydrogenase. We therefore examined whether these compounds restrict WNV replication in human neural progenitor cells.

Effective PHGDH inhibition in cell cultures has been reported at 20-40 µM for homoharringtonine (*45*) and 50 µM for DSF (*46*). However, treatment of ReNcell VM neural progenitor cells with 20 µM homoharringtonine caused profound cell death. A cytotoxicity assay revealed that homoharringtonine induces up to 80% cell death in neural progenitors at concentrations as low as 1.125 µM (Fig. 6C), whereas 50 µM DSF did not affect cell viability over 72 h of treatment (Fig. 6D). DSF was therefore selected for assessing its effect on WNV replication.

To assess the effect of DSF on WNV replication, ReNcell VM cells were infected with WNV_NSW2011_ at MOI=0.1 and then maintained in media supplemented with 50 µM DSF. Viral replication assay demonstrated that DSF treatment significantly reduced WNV replication in neural progenitor cells (Fig. 6E), with titres four orders of magnitude below vehicle control at 24 hpi. Titres in DSF-treated cells rose at later time points, possibly reflecting loss of drug activity over time, but remained on average 2 log_10_ lower than control with difference being statistically significant (Fig. 6E). This indicates that DSF efficiently inhibits WNV replication in human neural progenitor cells.

To assess whether the antiviral effect of disulfiram extends to multicellular three-dimensional neural tissue, we tested the drug in WNV-infected organoids. Human cerebral organoids were inoculated with 10 FFU of WNV_NSW2011_ per organoid and maintained in medium containing 50 µM DSF over a 5-day time course. DSF significantly reduced WNV replication compared with vehicle-treated controls at 4 and 5 dpi (Fig. 6F), indicating that the antiviral activity observed in monolayer culture is retained in a three-dimensional tissue context although delayed in time.

Collectively, these results identify disulfiram, a drug already approved for human use, as an inhibitor of WNV replication in both human neural progenitor cells and cerebral organoids. More broadly, our findings support pharmacological targeting of PHGDH as future host-directed antiviral strategy.

## DISCUSSION

In this study, we established a human cerebral organoid model of WNV_NSW2011_ infection and combined it with virus-inclusive scRNA-Seq to dissect the cellular basis of WNV neuropathogenesis. We found that WNV productively infects human cerebral organoids and exhibits a pronounced tropism for neural progenitors, astrocytes, and immature neurons, while largely sparing mature neurons. Astrocytes were identified as the principal innate immune responders to infection, whereas mature neurons mounted only a limited antiviral response that was likely associated with low expression of pattern-recognition receptors and interferon signaling components. The single-cell gene expression analysis also identified PHGDH, the rate-limiting enzyme of *de novo* serine biosynthesis, as a host dependency factor whose genetic depletion or pharmacological inhibition restricted WNV replication.

The human cerebral organoid model of WNV encephalitis has been recently independently established by Steffen *et al.* (*47*). Using an organoid system generated with the STEMdiff Cerebral Organoid Kit (an unguided protocol) and infected at DIV100 with a European lineage 1a WNV strain TOS-09 at a low dose (10^4^ FFU), the study demonstrated that human cerebral organoids support productive WNV infection without overt tissue destruction. They showed that viral antigen was localised to small foci in the outer cortical-like layers rich in neurons and astrocytes and documented a release of pro-inflammatory cytokines and chemokines in response to infection. A notable difference of our model concerns the reproducibility of infection. Steffen *et al.* reported pronounced heterogeneity in virus growth kinetics in individual organoids, with a bimodal distribution of viral replication peaks, maximum titers spanning six orders of magnitude (10^2^ to 10^8^ FFU/ml), and apparent viral clearance in approximately 37% of organoids (*47*). In contrast, the organoids used in our study displayed tight, convergent replication curves across individual organoids, inoculum doses, and maturation stages (Fig. 1A, B). The authors contribute heterogeneity to spontaneous differentiation of different amounts of the choroid plexus tissue by the mature age organoids used in the study (*47*). Greater consistency observed in our study likely reflects several factors including (i) our use of a directed, growth-factor-supplemented differentiation protocol, which produces organoids of more uniform cellular composition than the unguided cerebral organoid kit; (ii) the younger maturation stage of our organoids (DIV40 and DIV60 versus DIV100), at which tissue architecture is more homogeneous and choroid plexus formation is not observed; (iii) using Matrigel-free organoids that are more accessible to the viral particles present in the inoculum; and (iv) automated production and monitoring workflow. Importantly, both independently developed organoid models converged on the permissiveness of human cortical tissue for WNV infection and on the involvement of astrocytes in virus-host interactions in human neural tissue. This reinforces the validity of the cerebral organoid as a platform for studying WNV neural infection.

The astrocyte infection and innate immune reaction observed in our study are consistent with the earlier findings, that commonly reported astrocyte activation and astrogliosis in flavivirus-infected brain tissue (*48–50*). However, the resistance to infection of mature neurons contrasts with the previous findings that reported neuronal tropism of WNV in *in vivo* mouse models and cultured murine cells. In mouse CNS, WNV was reported to have a profound neuronal infection, with viral antigen detected in the neurons within multiple brain regions (*18, 51*). In *ex vivo* murine spinal cord slice cultures, by day 7 post infection the majority of neurons were infected and exhibited signs of apoptotic death (*50*). WNV was also shown to efficiently infect murine primary neurons (*15, 52, 53*), Neuro2a neuronal cell line and mouse embryonic stem cell (ES)-derived neurons (*18*). Human post-mortem studies of the fatal West Nile encephalitis cases (*54, 55*) reported presence of WNV antigen in neurons, although the neuronal identity of the cells was assigned solely based on cell morphology without immunodetection of the cell type markers and prevalence of neuronal vs glial infection hasn’t been elucidated. Steffen *et al.* used immunofluorescent staining of infected cerebral organoids to show that WNV envelope protein localized to outer cortical areas of the organoid that were rich in MAP2-positive neurons and GFAP-positive astrocytes (*47*). However, the definitive localisation of viral antigen with a cell expressing a specific cell-type marker was not demonstrated. The study also did not include markers of neural progenitor cells such as PAX6 and SOX2 or specific markers of neuronal maturity and therefore could not reveal infection of progenitors and immature vs mature neurons. In this study we demonstrated preferential tropism of WNV to neural progenitors/immature neural cells and astrocytes and poor tropism to mature neurons in cerebral organoids using two complementary approaches, scRNA-seq and immunostaining. The findings were validated using an alternative model of 2D human neural cell cultures infected with two different viral strains, confirming that the observed tropism is not due to genetics of the specific iPSC line used to generate organoids or NSW2011 viral strain. Notably, a recent study that used iPSC-derived monocultures of human neurons, astrocytes and neural progenitor cells also reported high WNV tropism to glial cells and a remarkable resistance of the mature neurons (*56*). Collectively the discrepancies between the studies that utilised human and rodent infection systems point towards a potential species-specific nature of WNV tropism and highlight the importance of further research in biologically relevant human models.

Our scRNA-Seq analysis identified astrocytes as the dominant innate immune effector against WNV in human cortical tissue. Astrocytes were the principal source of type I and type III interferons and also mounted a robust ISG response, acting as the main hub of inferred interferon signalling. This is consistent with evidence that astrocytes rapidly restrict neurotropic flaviviruses through a cell-intrinsic interferon response and maintain elevated basal expression of antiviral effectors (*48*). In contrast, mature neurons showed limited ISG induction. By profiling the upstream signalling machinery, we found that this hyporesponsiveness has a clear molecular correlate with low levels of viral RNA sensors and key interferon receptor signalling components in the mature neurons. This immunological deficiency would be expected to make neurons vulnerable to infection due to the lack of protection provided by ISG expression, which was not evident. Therefore, we hypothesised that poor susceptibility of mature neurons to the virus likely results from the lack of expression of the host factor(s) required for viral propagation.

Differential expression analysis between WNV-infected and virus-negative cells revealed high expression of serine biosynthesis and one-carbon metabolism PHGDH, SHMT2 and MTHFD2 as a transcriptomic signature of WNV-permissive cells. These genes were not induced by infection, which indicates that they mark a pre-existing permissive cell state rather than a virus-induced response. PHGDH has previously been linked to viral replication, although its reported role is contradictory. Li *et al.* showed that classical swine fever virus (CSFV) promotes PHGDH degradation, reducing serine biosynthesis, MAVS-IRF3 signalling and IFN-β production, enhancing viral replication and proposing an antiviral role for PHGDH (*57*). In contrast, Shen *et al.* found that Sendai virus (SeV) and vesicular stomatitis virus (VSV) down-regulate *Phgdh*, *Shmt2*, and *Mthfd2* genes in murine macrophages and showed that serine metabolism suppresses type I IFN responses, assuming a pro-viral role of serine pathway (*58*). Although both studies describe suppression of serine metabolism during infection, they assign opposing consequences to this response. Our findings suggest a different relationship between the virus and serine pathways in human neural context. We observed no significant change in PHGDH expression following WNV infection in human neural progenitor cells, providing evidence that WNV doesn’t actively remodel this pathway. Instead, elevated basal expression of serine synthesis and one-carbon metabolism genes was associated with cells supporting viral replication. We propose that this observation is best understood not as a virus-specific interaction but as a reflection of the metabolic state that WNV requires. PHGDH catalyses the rate-limiting, committed step of *de novo* serine synthesis (*59, 60*). Serine is a central biosynthetic hub, which, through SHMT2 and MTHFD2, fuels the folate-mediated one-carbon cycle that supplies purine and thymidine synthesis. It also provides precursors for phospholipid and sphingolipid production, regenerates the methyl donor S-adenosylmethionine, and contributes to NADPH-dependent redox balance (*61*). Each of these outputs is heavily utilised by flavivirus replication. As all flaviviruses, WNV amplifies its positive-strand RNA genome to high copy number, which has a high demand on nucleotides. It also remodels the endoplasmic reticulum into lipid-rich replication organelles (*62–64*), and methylates its RNA cap at the 2′-O position through the NS5 methyltransferase to evade IFIT-mediated restriction (*65*), which requires lipids and the source of one-carbon groups respectively. A high-flux serine and one-carbon pathway therefore represent a single coordinated metabolic module which supplies the nucleotides, membrane lipids, and methyl groups required for virus replication.

This interpretation also unifies the two principal findings of our study. *De novo* serine synthesis is a hallmark of proliferative state and anabolic metabolism. Therefore, PHGDH is usually highly expressed in dividing and cancer cells (*59, 60*). In the cerebral organoids, the cells with the highest anabolic rate and therefore the serine-synthesis activity are the cycling progenitors and radial glia, whereas post-mitotic mature neurons have low demand for biosynthetic processes and correspondingly low serine pathway activity, relying on serine produced by astrocytes for their limited needs (*66*). WNV tropism for progenitors and astroglia over mature neurons, and the dependency of viral replication on PHGDH, are thus two manifestations of the same underlying phenomenon. The virus exploits the biosynthetically primed metabolic state of immature dividing neural cells. Consistent with this, TRIB3 and EIF4EBP1, the proteins most strongly associated with infected cells alongside PHGDH (Fig. 5A), are the effectors of the ATF4-driven stress response, which also upregulates serine-synthesis and one-carbon enzymes (*67*).

We further demonstrated that RNAi knock-down and pharmacological inhibition of PHGDH with CBR-5884 and DSF reduced viral replication in permissive neural cells. This nominates *de novo* serine synthesis as a target for development of host-directed antiviral therapies against WNV neuro-infection. Such interventions that target cellular processes co-opted by the virus rather than viral proteins, offer a higher genetic barrier to resistance and can potentially be applicable to related viruses. The efficiency of the FDA-approved drug DSF against WNV in neural progenitor culture and human cerebral organoids provides the proof-of-concept experiment that the available safe drugs can be potentially repurposed to target PHGDH and reduce WNV replication in human neural tissue. We noticed that the effect of DSF in the organoids emerged later than in two-dimensional culture (Fig. 6E,F), which is consistent with the requirement for the drug to traverse organoid tissue into which small molecules penetrate over a timescale of hours and to a depth of only a few hundred micrometers (*68*). The lower magnitude of the antiviral effect of the drug at the late time points is likely explained by the limited chemical stability of DSF in aqueous, protein-containing media as DSF is rapidly reduced to diethyldithiocarbamate (DDC) by free thiols and by serum albumin (*69*), with a reported half-life of under 2 min in serum (*70*). Although these limitations, together with relatively high concentration of the compound required for PHGDH inhibition (50 μM) (*46*), will likely preclude the clinical use of DSF itself against WNV, our results establish PHGDH as a druggable host dependency factor in human neural tissue and provide a rationale for developing more potent and metabolically stable inhibitors of *de novo* serine synthesis or other approaches for serine pathway suppression. PHGDH has been pursued intensively as an oncology target, and small-molecule inhibitors, together with the concept of dietary serine and glycine restriction, are under active investigation in the field of cancer therapy (*59–61*). This provides a future pipeline of interventions and compounds that can be potentially repurposed to limit WNV replication in the CNS.

## MATERIALS AND METHODS

### Cell culture

The human astroglial cell line SVG p12 (ATCC, CRL-8621) and the human microglial cell line HMC3 (ATCC, CRL-3304) were maintained in Eagle’s Minimum Essential Medium (EMEM) (ATCC, USA) supplemented with 10% fetal bovine serum (FBS). African green monkey kidney cell Vero cells (ATCC, CCL-81) were cultured in Dulbecco’s Modified Eagle Medium (DMEM) with 10% FBS. The immortalised human ventral mesencephalic neural progenitor cell line RENcell VM (Merck Millipore, SCC008) was propagated as an adherent monolayer on Matrigel-coated cell culture vessels. Culture surfaces were coated with hESC-qualified Matrigel Matrix (Corning, USA, #354277) diluted to 1% (v/v) in Dulbecco’s phosphate-buffered saline (DPBS) without calcium or magnesium (Thermo Fisher Scientific, USA) using 5 ml per T75 flask or 1 ml per well of a 6-well plate, followed by incubation for 1 hour at 37°C. Residual coating solution was aspirated immediately before use. Cells were propagated in DMEM/F12 supplemented with GlutaMAX (Thermo Fisher Scientific, USA, #10565018), 2% (v/v) B-27 Supplement (Thermo Fisher Scientific, USA, #17504044), 20 ng/ml recombinant human EGF (STEMCELL Technologies, USA, #78006.2), and 20 ng/ml recombinant human bFGF (STEMCELL Technologies, USA, 78003). All mammalian cell cultures were maintained at 37°C in humidified atmosphere of 95% air and 5% CO. C6/36 Aedes albopictus cells (ATCC, CRL-1660) were maintained in Roswell Park Memorial Institute (RPMI 1640) medium supplemented with 10% FBS at 28 °C. Unless otherwise stated all cell culture reagents were from Gibco, USA. All cells were confirmed free of mycoplasma.

### Generation of hiPSC-derived human cerebral organoids

Cerebral organoids were generated from the human induced pluripotent stem cell (hiPSC) line WTC11 (Coriell Institute for Medical Research, GM25256) following a modified version of the protocol by Anastasaki et al (*34*) and were matured for 40 or 60 days. Unless otherwise stated, cultures were maintained at 37°C in a humidified 5% CO incubator, and organoid formation and maintenance were carried out using automated liquid handling. WTC11 hiPSCs were maintained under feeder-free conditions on tissue-culture plates coated with hESC-qualified Matrigel Matrix (Corning, USA, #354277) in mTeSR Plus medium (STEMCELL Technologies, USA, #100-0276) and were passaged as aggregates using ReLeSR (STEMCELL Technologies, USA #100-0483). Cells were monitored for morphology, confluency and the absence of spontaneous differentiation, and were expanded to the required confluency before organoid generation.

For embryoid body formation and neural induction, the hiPSCs were dissociated to a single-cell suspension with Accutase Cell Dissociation Reagent (STEMCELL Technologies, USA, #07920), and 20,000 cells were aggregated per well in U-bottom ultra-low-attachment 96-well plates (prepared with Anti-adherence Rinsing Solution; STEMCELL Technologies, USA, #07010) to form uniform embryoid bodies (EBs). EBs were cultured in Neural Induction Medium (NIM) consisting of Neurobasal Medium (Thermo Fisher Scientific, USA, #21103-049) supplemented with 1x GlutaMAX (Thermo Fisher Scientific, USA), 1x penicillin/streptomycin (Thermo Fisher Scientific, USA), 1x N-2 Supplement (Thermo Fisher Scientific, USA, 317502048), 2x B-27 Supplement (Thermo Fisher Scientific, USA, #17504001), 1.25 µM dorsomorphin (STEMCELL Technologies, USA, #72102), 3 µM CHIR99021 (STEMCELL Technologies, USA, #72054), 3 µM SB431542 (STEMCELL Technologies, USA, #72232) and 10 ng/ml recombinant human LIF (STEMCELL Technologies, USA, #78055). For the first four days, NIM was additionally supplemented with 10 µM Y-27632 ROCK inhibitor (STEMCELL Technologies, USA, #72308) to enhance cell survival and aggregate consistency. Complete medium changes were performed on days 5 and 8 in the absence of ROCK inhibitor.

On day 10, EBs were transferred to low-attachment 24-well plates for long-term free-floating culture and were maintained on an orbital shaker in Cerebral Brain Organoid Medium (CBOM). CBOM comprised Neurobasal Medium (Thermo Fisher Scientific, USA, #21103-049) and DMEM/F-12 (Thermo Fisher Scientific, 11320-033) in 1:1 ratio supplemented with 1x GlutaMAX (Thermo Fisher Scientific, USA), 1x antibiotic-antimycotic (Thermo Fisher Scientific, USA, #15240062), 1x N-2 Supplement (Thermo Fisher Scientific, USA, #17502048), 2x B-27 Supplement (Thermo Fisher Scientific, USA, #17504001), 100 U/ml human recombinant insulin (Thermo Fisher Scientific, USA, #12585014), 200 nM 2-mercaptoethanol (Sigma-Aldrich, USA, #M6250), 0.5x MEM non-essential amino acids (Thermo Fisher Scientific, USA, #11140-050), and 20 ng/ml each of recombinant human BDNF (STEMCELL Technologies, USA, #78133.1), GDNF (STEMCELL Technologies, USA, #78139.1) and FGF2 (STEMCELL Technologies, USA, #78134.1). Organoids were matured to day 40 and day 60 with medium refreshed three times per week and were routinely monitored by whole-well brightfield imaging on a Celigo image cytometer (Nexcelom) before each medium change. The organoids were produced by Australian Organoid Facility (The University of Queensland, Brisbane, QLD).

### Neuron Differentiation

Differentiation of RENcell VM neural progenitors cells into neurons was carried out on sequentially coated poly-L-ornithine/laminin attachment surfaces. Cell culture vessels were first coated with undiluted poly-L-ornithine solution (Merck, USA, #P4957), applying 5 ml per T75 flask or 500 μl per well of a 6-well plate and incubated overnight at 37°C. The poly-L-ornithine was aspirated and the surface was washed once with DPBS (Thermo Fisher Scientific, USA). Vessels were then coated with 1% (v/v) laminin, prepared by diluting mouse natural laminin (Thermo Fisher Scientific, USA, #23017015) to 1% in DPBS and incubated overnight at 37°C. The laminin solution was aspirated immediately before seeding, and the surface was not washed. RENcell VM cells were seeded onto the coated surface at a density of 2.5-5×10 cells/cm² in Expansion Medium and allowed to attach for 48 hours at 37°C/5% CO. To initiate differentiation, expansion medium was replaced with Neural Differentiation Medium consisting of Neurobasal Medium (Thermo Fisher Scientific, USA, #21103049) supplemented with 2% (v/v) B-27 Supplement (Thermo Fisher Scientific, USA, #17504044), 1% (v/v) GlutaMAX Supplement (Thermo Fisher Scientific, USA, 35050061), and 1% (v/v) Insulin-Transferrin-Selenium-X (ITS-X) (Thermo Fisher Scientific, USA, #51500056). Growth factors (EGF and bFGF) were withdrawn at this step. Thereafter, cells were gently washed and refed with fresh differentiation medium every 2–4 days, and differentiation was allowed to proceed for additional 12 days until a mature neuronal phenotype was attained.

### Viruses and infection

WNV_NSW2011_, the Australian Kunjin lineage 1 strain of WNV isolated during the 2011 equine encephalitis outbreak (*5*) was obtained from Prof Peter Kirkland (Elizabeth Macarthur Agricultural Institute, NSW, Australia) and then recreated using Circular Polymerase Extension Reaction (CPER) (*71*). The North American strain WNV_NY99_ was obtained from Prof Roy Hall (University of Queensland, QLD, Australia) and then recreated by CPER (*72*). The viral genomes were sequenced and confirmed to match GeneBank reference sequences JN887352 for WNV_NSW2011_ and NC_009942 for WNV_NY99_. Viruses were propagated in Vero cells and used at passage 2.

For organoid infections, individual organoids were inoculated with 10^4^ or 10^5^ FFU of WNV_NSW2011_ in 500 ul of Cerebral Brain Organoid Medium (CBOM) and incubated overnight at 37 °C and 5% CO_2_. The inoculum was then removed, organoids were washed twice with serum-free F12K medium, and fresh CBOM was added. Two-dimensional cell cultures were infected with WNV_NSW2011_ or WNV_NY99_ at indicated multiplicity of infection (MOI) for 1h at 37 °C in 6-well plates using 50 μl of inoculum per cm^2^ of growth area. After inoculum was removed, cells were washed twice with 2 ml/well PBS and then maintained in the appropriate culture medium. For HMC3 and SVG p12 cells, the reduced serum medium (2% FBS) was used to prevent overgrowth.

### Virus titration

WNV titres were determined using an immunofluorescent foci-forming assay (*73*). Culture fluid samples were used to prepare 10-fold serial dilutions in RPMI 1640 supplemented with 2% FBS, and 25 μl of each dilution was added to 10^5^ C6/36 cells/well seeded in 96-well plates. Following adsorption for 1 h at 28 °C, cells were overlaid with 175 μl of overlay medium and incubated at 28 °C with 5% CO. The overlay consisted of equal volumes of 2x M199 medium (Gibco, USA) containing 5% FCS, 2.2 g/L NaHCO (Merk, USA), and 2% carboxymethyl cellulose (Merk, USA).At 3 days post-infection, cells were fixed with ice-cold PBS-buffered 80% acetone (100 μl per well) for 20 min at −20 °C, air-dried, and blocked for 30 min with 50 μl/well Pierce Clear Milk blocking solution (Thermo Fisher Scientific, USA). Viral foci were detected by incubation with the mouse monoclonal antibody 4G2 specific for flavivirus envelope protein (*74*) diluted 1:100 (50 μl/well), followed by goat anti-mouse IRDye 800CW secondary antibody (LI-COR, USA) diluted 1:2000 (50 μl/well). Antibodies were prepared in Clear Milk blocking buffer, and incubations were carried out for 1 h at 37 °C. Plates were washed five times with PBS containing 0.05% Tween 20 (PBST) after each antibody incubation. Plates were air-dried and fluorescent virus replication foci were visualised using Odyssey Classic Imaging System (LI-COR, USA) (42 μm; medium resolution; 3.0 mm). Foci were counted using Image Studio Lite software (LI-COR, USA). Infectious titres were calculated from the dilution factor and reported as focus-forming units per milliliter (FFU/ml). Whole-mount immunofluorescent analysis of cerebral organoids. Organoids were rinsed in DPBS (Thermo Fisher Scientific, USA) and fixed in 4% paraformaldehyde freshly diluted from a 16% (w/v) stock (Thermo Fisher

Scientific, #043368.9M) in DPBS for 30 min at room temperature, followed by three washes in DPBS. Organoids were transferred to 1.5 ml tubes and permeabilised/blocked overnight at room temperature on a rocker in blocking buffer comprising 6% (w/v) bovine serum albumin (from Blocker BSA 10% in PBS; Thermo Fisher Scientific, USA, #37525) and 0.2% (v/v) Triton X-100 (Sigma-Aldrich, x100) in DPBS. Organoids were then incubated with primary antibodies diluted in blocking buffer overnight at room temperature on a rocker. The primary antibodies used were rabbit anti-GFAP (Abcam, UK, #ab68428), rabbit anti-MAP2 (Abcam, UK, #ab11267), rabbit anti-β III tubulin/TUJ1 (Abcam, UK, #ab78078), rabbit anti-neurofilament (Cell Signaling Technology, USA, #2837S), and rabbit anti-PAX6 (Invitrogen, USA, #42-6600). Organoids were then washed three times, 10 min each, in 0.2% (v/v) Triton X-100 in DPBS (PBS-T) on a rocker and incubated with secondary antibodies diluted in blocking buffer overnight at 4°C on a rocker. Secondary antibodies were PhenoVue Fluor 488 goat anti-rabbit IgG (Revvity, USA, #2GXRB488C1) and PhenoVue Fluor 647 goat anti-mouse IgG (Revvity, USA, 2GXM647C1). Following a single PBS-T wash, nuclei were counterstained with DAPI (Invitrogen, USA, #D1306) diluted in blocking buffer for 1 h at room temperature on a rocker, and then washed three times, 10 min each, in PBS-T. Stained organoids were mounted in glass-bottom plates and imaged on an Opera Phenix Plus High-Content Imaging System (Revvity, USA) using a 20x water-immersion objective. Images are presented as maximum-intensity projections of 111 optical planes acquired at 5 µm z-spacing.

### Organoid dissociation

Organoids were dissociated into single-cell suspensions using the Papain Dissociation System (Worthington Biochemical Corporation, USA) following the manufacturer’s guidelines with modifications optimized for organoid tissue. For each condition, 10 organoids were pooled to yield approximately 2×10^6^ cells. The papain solution was prepared by reconstituting the lyophilized papain vial in 5 ml of Earle’s Balanced Salt Solution (EBSS) and incubating at 37 °C for approximately 10 min until fully dissolved. In parallel, DNase was reconstituted in 500 µl of EBSS and 250 µl of the resulting DNase solution was added to the dissolved papain to generate the working dissociation mix. For each condition, 10 organoids were pooled in a well of a 6-well plate, washed once with PBS, and then cut into small pieces using a sterile straight scalpel, taking care to avoid over-chopping. The papain-DNase mix (1 ml) was added to the minced tissue, and the plate was incubated at 37 °C, 5% CO_2_ for up to 40 min with gentle orbital agitation. During incubation, samples were subjected to gentle pipetting 4-5 times every 5 min. Incubation was terminated upon complete dissociation of the organoids with no tissue clumps visible. The resulting cell suspension was filtered through a 40 µm EasyStainer cell strainer (Greiner Bio-One, Austria) placed on a 50 ml conical tube to isolate single cells. The strainer was first pre-wetted with 1 ml of Cerebral Brain Organoid Medium (CBOM), after which the dissociated cells were collected and passed through the strainer in two sequential 5 ml washes of medium. The filtered suspension was transferred to a 15 ml conical tube and centrifuged at 300 x g for 5 min at room temperature. The cell pellet was resuspended in 1 ml of CBOM, cell numbers and viability were determined by trypan blue exclusion staining using Countess 3 Cell Counter (Thermo Fisher Scientific, USA). Cells were used immediately for formaldehyde fixation.

### Virus-inclusive single-cell RNA sequencing (scRNA-Seq) and bioinformatics analysis

Cells from the dissociated organoids (10^6^ cells per sample) were fixed using Chromium Next GEM Single Cell Fixed RNA Sample Preparation Kit (10x Genomics, USA) according to the manufacturer’s instructions and counted on Countess 3 Cell Counter (Thermo Fisher Scientific, USA) using trypan blue staining. Fixed cells were used for single cell library preparation with Chromium Fixed RNA Kit, Human Transcriptome 4 rxns x 4 BC (10x Genomics, USA). At the probe hybridisation step, 10x Genomics human transcriptome probes were supplemented with probes complementary to WNV genome (Supplementary Table S1) according to the manufacturer’s instructions. Cells (10,000 per sample) were partitioned using Chromium Next GEM Chip Q Single Cell chip (10x Genomics, USA) and Chromium X instrument (10x Genomics, USA). Libraries were barcoded using Dual Index Kit TS Set A (10x Genomics, USA), analysed on Agilent TapeStation 4200 and sequenced on Illumina NovaSeq 6000 Sequencing System. Initial data pre-processing was performed using Cell Ranger v9.0.1 (10x Genomics) using custom-built genome to which viral genomic sequence was added as an extra chromosome. Single-cell data was then analysed in R (v4.5.1) using Seurat (v5.5.0) following the standard workflow. Filtered gene-barcode matrices were imported and used to construct a Seurat object. Low-quality cells and putative doublets were excluded on the basis of the number of detected genes (nFeature_RNA), total unique molecular identifier counts (nCount_RNA) and the percentage of mitochondrial reads, the latter calculated with PercentageFeatureSet using the “^MT-” gene prefix. Cells with fewer than 300 or more than detected genes, or with more than 10^5^ UMIs or 5% mitochondrial content, were removed. Filtered counts were normalised using the global-scaling method and the 2,000 most variable genes were identified with FindVariableFeatures. Data were scaled and centred with ScaleData with mitochondrial content regressed out, then linear dimensionality reduction was performed by principal component analysis (RunPCA) on the variable genes. The dimensionality of the dataset was assessed by elbow plot, and the first 13 principal components were retained for downstream analysis. A shared-nearest-neighbor graph was performed with FindNeighbors using the selected principal components and cells were clustered with FindClusters (Louvain algorithm) at a resolution of 0.4. Clusters were visualised in two dimensions by uniform manifold approximation and projection (RunUMAP) over the same principal components. Cluster marker genes were identified with FindAllMarkers using the Wilcoxon rank-sum test. Differentially expressed genes were considered significant at an adjusted P value < 0.05 (Bonferroni correction). Clusters were annotated to cell types on the basis of the expression of the canonical marker genes.

The WNV genome was included in the reference as an additional feature (“WNV”), and viral loads were quantified per cell as WNV unique molecular identifier (UMI) counts. Cells were classified as infected if WNV counts were at least 2 UMI. The proportion of infected cells and the mean WNV UMI per cell were calculated for each cluster at each timepoint. Intercellular signalling was inferred using a custom ligand-receptor scoring approach based on a curated ligand-receptor database. For each sender-receiver cluster pair, a communication score was calculated as the product of mean ligand expression in the sender and mean receptor expression in the receiver (AverageExpression), summed across all pairs; scores were aggregated per signalling pathway. Communication was visualised as sender-by-receiver heatmaps. Differential gene expression analyses were performed using the Wilcoxon rank-sum test. Functional over-representation analysis of significantly up-regulated genes (per cluster and contrast) was performed using clusterProfiler v4.16.0 after mapping of gene symbols to Entrez IDs with org.Hs.eg.db v3.21.0. Testing was performed for Gene Ontology Biological Process terms (enrichGO) and KEGG pathways (enrichKEGG, organism “hsa”) with Benjamini-Hochberg correction (P < 0.05; GO q < 0.1). Gene sets with fewer than 10 mapped genes were excluded. Fold enrichment was calculated as the ratio of gene ratio to background ratio, and results were displayed as dot plots. Data visualisations were performed using ggplot2 v4.0.3 or native Seurat plotting capabilities.

### Immuno-histological staining of the cerebral organoids

Organoids were fixed in freshly prepared 4% paraformaldehyde solution in PBS overnight at 4°C, washed with PBS and embedded in 2.5% low-melting-point agarose (Promega, USA). Tissues were then dehydrated, processed, embedded in paraffin, and sectioned at 4 µm. Sections were subjected to dual-marker immunofluorescence in which flaviviral envelope antigen was co-detected with neural cell-type markers. Heat-induced epitope retrieval was performed in a Biocare Medical DIVA antigen retrieval buffer at 100°C for 20 min for NeuN and S100B. For doublecortin and Sox2, retrieval was performed in DIVA antigen retrieval buffer (Biocare Medical, DV2004) at 110°C for 10 min. Sections were then blocked in Background Sniper (Biocare Medical, BS966) supplemented with 2% (w/v) BSA and probed overnight with anti-flavivirus group antigen (envelope) monoclonal antibody 4G2 (*74*) diluted 1:20 in Da Vinci Green antibody diluent (Biocare Medical, PD900) followed by 1h incubation with Alexa Fluor 488-conjugated anti-mouse IgG (Invitrogen, USA; 1:500). Sections were then counter-stained for cell type markers by probing with either rabbit monoclonal antibody against doublecortin (#ab207175 Abcam, UK; 1:250), rabbit anti-Sox2 (#ab92494, Abcam, UK; 1:100), rabbit anti-NeuN (#ab17748 Abcam, UK; 1:600) or S100B (#NCL-L-S100p Leica Biosystems/Novocastra, UK; 1:150) for 1h followed by an Alexa Fluor 555-conjugated anti-rabbit IgG secondary antibody (Invitrogen, USA; 1:500). Nuclei were counterstained with 4′,6-diamidino-2-phenylindole (DAPI). Slides were imaged on Axioscan 7 high-performance automated slide scanner (Zeiss, Germany) using 20x objective. Images were analysed using Fiji distribution of ImageJ2 software.

### Immunofluorescence assay (IFA) in cultured cells

RENcell VM neural progenitor cells were seeded on poly-L-ornithine/laminin-coated glass coverslips contained in a 24-well plate and neuron differentiation was induced by growth factors withdrawal as described above. At 6, 12 and 14 days of differentiation cells were washed once with DPBS and fixed with ice-cold 100% methanol (500 μl per well) at −20°C for 20 min. Fixed cells were rinsed twice with 500 μl of Tris-buffered saline (TBS) and incubated with 500 μl of 50 mM glycine for 30 min at room temperature to quench autofluorescence. Cells were then washed three times for 5 min with 500 μl of TBS, blocked for 1 hour at room temperature with 500 μl of 1% bovine serum albumin in TBS, and incubated at 37°C for 1 hour with 250 μl of primary antibodies diluted in blocking solution. The mixture of antibodies contained rat monoclonal antibody against NeuN (#ab279297, Abcam, UK) diluted 1:1000 and mouse monoclonal antibody against MAP2 (#M9942, Merk, USA) diluted 1:1000. Cells were then washed three times for 5 min with TBS (500 μl per well) and incubated at 37°C for 1 hour with 250 μl of a mixture containing donkey anti-rat immunoglobulin G (IgG) (H+L) highly cross-adsorbed secondary antibody conjugated with Alexa Fluor 594 (Invitrogen, USA) and goat anti-mouse IgG (H+L) highly cross-adsorbed secondary antibody conjugated with Alexa Fluor 648 (Invitrogen, USA), each diluted 1:500 in blocking solution. After the incubation, cells were washed three times for 5 min with 500 μl of TBS per well, and coverslips were mounted using ProLong Gold Antifade reagent with 4′,6-diamidino-2-phenylindole (DAPI) (Invitrogen, USA). Imaging was performed using Leica Stellaris 5 Confocal Microscope.

### RNA interference

ReNcell VM neural progenitors were seeded in 24-well plates and transfected with 50nM siGENOME Human PHGDH siRNA SMARTpool (Horizon Discovery, UK, #M-009518-01-0005) or siGENOME Non-targeting Control siRNA (Horizon Discovery, UK, #D-001210-01). Transfection was performed using Dharmafect 2 transfection reagent (Horizon Discovery, UK). At 24 hours post-transfection, cells were infected with WNV_NSW2011_ at an MOI=0.1 for 1h. Cells were washed three times with DPBS and then maintained in 1m media per well. Culture fluid samples (50 μl) were collected for titration every 24h.

### Western Blotting

Cells were lysed in 150 μl per well of Bolt LDS sample buffer (Thermo Fisher Scientific, USA) supplemented with Bolt sample reducing agent (Thermo Fisher Scientific, USA) and protease inhibitor cocktail (Sigma-Aldrich, USA). Lysates were sonicated using a Branson Sonifier SLPt (Marshall Scientific) at 10% output for 15 s with a pulse on for 3 s and off for 1 s. Lysates were then incubated at 95°C for 5 min, cooled on ice, and loaded into the Bolt 4-12% bis-tris gels (Thermo Fisher Scientific, USA). Electrophoresis was performed for 30 min at 200 V in Bolt MES SDS running buffer (Thermo Fisher Scientific, USA). Proteins were then electroblotted onto a nitrocellulose membrane using iBlot3 Dry Blotting System (Thermo Fisher Scientific, USA) and iBlot 3 Transfer Stack (Thermo Fisher Scientific, USA) at 25V for 6 min. Membranes were blocked in 5% skim milk in Tris-buffered saline pH=7.4 with 0.05% Tween 20 (TBS-T) for 1 hour, incubated with primary antibodies overnight at 4°C, then washed four times for 5 min with TBS-T, incubated with horseradish peroxidase (HRP)-conjugated secondary antibodies for 1 hour at room temperature, and washed four times for 5 min with TBS-T. Detection of HRP activity was performed using SuperSignal West Pico PLUS chemiluminescence substrate (Thermo Fisher Scientific, USA), and signal was visualized using iBright CL1500 gel documentation instrument (Thermo Fisher Scientific, USA). The primary antibodies were rabbit monoclonal anti-PHGDH (D8F30) (Cell Signalling Technologies, USA, #66350) diluted 1:1,000 and mouse monoclonal anti-GAPGH (GAPDH-71.1) (Merk, USA, #G8795) diluted 1:10,000. The secondary antibodies were anti-rabbit IgG, HRP-linked (Cell Signalling Technologies, USA, #7074) and anti-mouse IgG, HRP-linked (Cell Signalling Technologies, USA, #7076), both diluted 1:2,500. All antibodies were diluted in 5% skim milk in TBS-T.

### Cytotoxicity Assay

Cytotoxicity of the chemical compounds was assessed using the CellTiter 96 AQueous One Solution Cell Proliferation Assay (Promega, USA) according to the manufacturer’s instructions. ReNcell VM neural progenitor cells were seeded into Matrigel-coated 96-well plates in Expansion Medium and allowed to attach for 24 h at 37°C in a humidified 5% CO incubator. Compounds were prepared as concentrated stocks in dimethyl sulfoxide (DMSO) and diluted in Expansion Medium to their respective starting concentrations, which were then serially diluted two-fold across the plate. The starting concentrations were 30 µM for CBR-5884 (#24471 New England Biolabs, USA), 50 µM for disulfiram (#D2950000 Merk, USA), and 10 µM for homoharringtonine (#SML1091 Merk, USA). Cells were treated in three replicate wells per concentration, with vehicle-only (DMSO) and medium-only wells included as negative controls. Treated plates were incubated for 24-72h at 37°C, 5% CO. Following incubation, 20 µl of CellTiter 96 AQueous One Solution reagent was added to each well containing 100 µl medium, and plates were incubated for 1-2h at 37°C, 5% CO protected from light, then the reaction was stopped by addition of 25 µl of 10% SDS per well. Absorbance was measured at 490 nm on a POLARstar OPTIMA microplate reader (BMG Labtech, Germany). Cell viability was expressed as a percentage of the untreated control.

### Compound treatment

RENcell VM neural progenitor cells were seeded into Matrigel-coated 96-well plates in Expansion Medium and allowed to attach for 24 h at 37°C in a humidified 5% CO incubator. Cells were then pre-treated with the PHGDH inhibitor CBR-5884 (#24471, New England Biolabs, USA) at 30 µM in Expansion Medium, or with an equivalent volume of DMSO-containing medial as a vehicle control, for 24 hours before infection. Cells were then infected with WNV_NSW2011_ at an MOI=0.1 for 1h, washed twice with 2ml/well of DPBS and then maintained in Expansion Medium containing 30 µM CBR-5884 or vehicle for 72 h. For DSF treatment, cells were maintained in Expansion Medium without the drug until virus inoculation, which was performed as for CBR-5884. After removal of inoculum and two washes with DPBS cells were maintained in Expansion Medium containing 50 µM DSF or matching volume of DMSO. Culture supernatants were collected at 24, 48, and 72 hpi for titration by immunofluorescent focus-forming assay.

Cerebral organoids were inoculated with 10^5^ FFU of WNV_NSW2011_ overnight, washed three times with F12K medium and then maintained in 1 ml/well of CBOM containing 50 µM DSF or matching volume of DMSO. Culture supernatants (50 µl) were sampled daily at 0-5 dpi and analysed by immunofluorescent focus-forming assay.

### Statistical analysis

Statistical analyses were performed using GraphPad Prism v.10.6.1. The statistical tests and sample sizes are indicated in figure legends and/or results section. A *P* value < 0.05 was considered statistically significant.

## Acknowledgments

The authors acknowledge the facilities and the scientific and technical assistance of The AUSTRALIAN ORGANOID FACILITY (AOF) at The University of Queensland. The AUSTRALIAN ORGANOID FACILITY (AOF) is supported by Phenomics Australia (PA). PA is supported by the Australian Government through the National Collaborative Research Infrastructure Strategy (NCRIS) program. The authors acknowledge the support of the Histology Facility at QIMR Berghofer Medical Research Institute for expert technical assistance with tissue processing, sectioning, staining, and slide preparation. We acknowledge the support of QIMR Berghofer microscopy facility in imaging and microscopy and University of Queensland Genomics Facility in single cell sequencing.

## Author contributions

Conceptualization: AS

Methodology: AS, ML, FS, NG, EW, MS, ALC, DJR

Investigation: ML, LDH, ZZ, FS, JXJC, WR, NQ

Visualization: AS

Funding acquisition: AS, AAK

Project administration: AS

Supervision: AS, AAK, ML, EW, NG

Writing – original draft: AS, ML

Writing – review & editing: AS, AAK, ML

