## Supplementary figures and tables for "Single-cell analysis of West Nile virus infection in brain organoids identified a serine synthesis enzyme as a determinant of viral neural tropism and a druggable antiviral target"

**Supplementary materials**

**Supplementary table S1. Sequences of the WNV-specific probes used for virus-inclusive scRNA-Seq.**

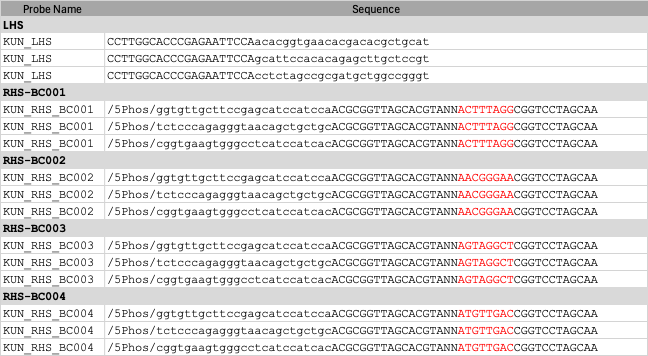

Constant probe segments are capitalised, pooling barcodes are shown in red.

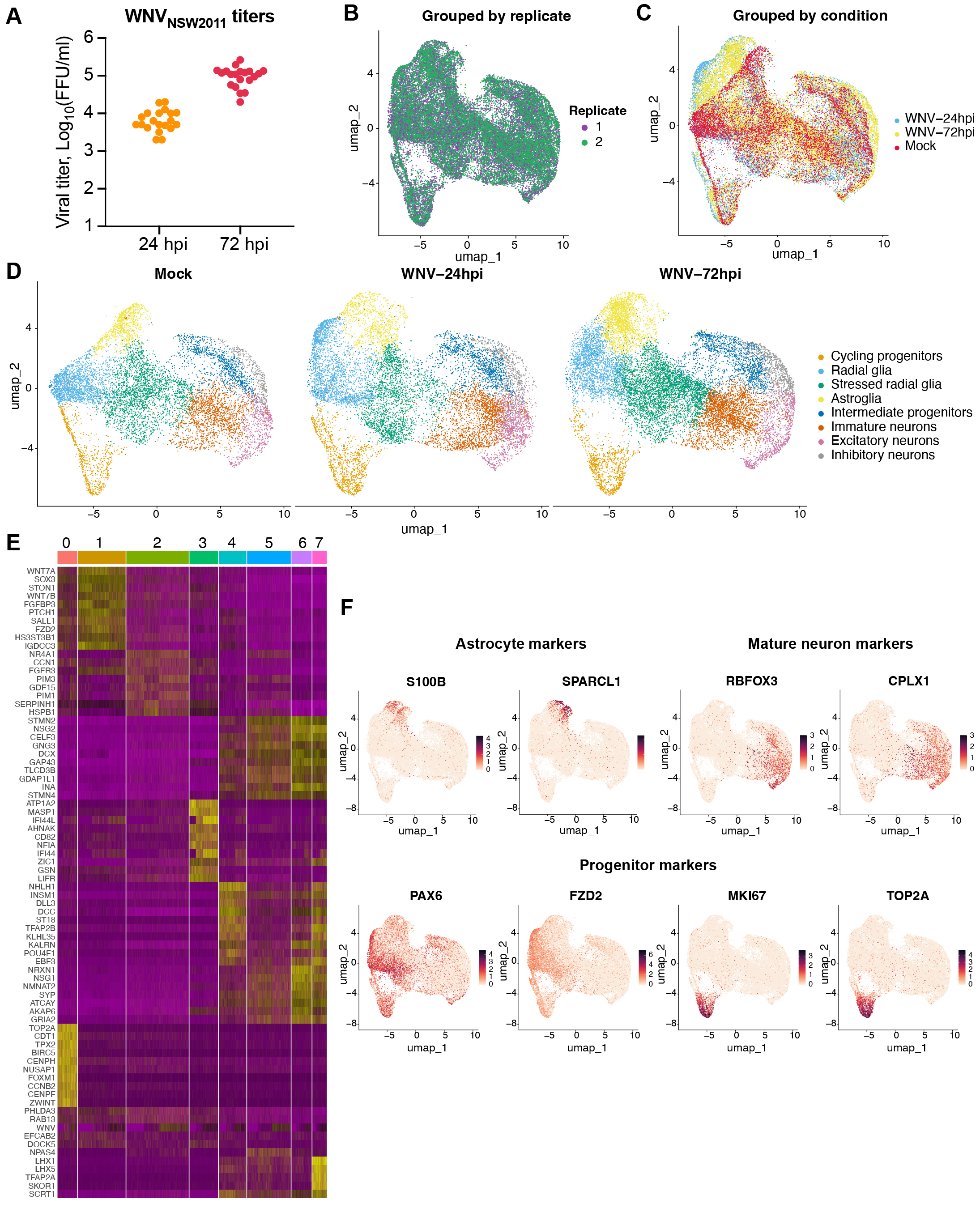

**Fig. S1. Cellular composition of mock- and WNV-infected human cerebral organoids identified by single cell RNA-Seq analysis. (A)** Viral titers in culture supernatants from WNV_NSW2011_-infected cerebral organoids used in scRNA-Seq experiment. Titers were determined by immunofluorescent focus-forming assay on C6/36 cells. Each point represents a measurement from an individual organoid; horizontal lines indicate the means. **(B)** Uniform manifold approximation and projection (**UMAP**) visualisation of single-cell transcriptomes of cerebral organoids with colouring by biological replicate. **(C)** UMAP visualisation of the same dataset coloured by experimental condition (mock-infected organoids, WNV-infected organoids at 24 hpi, and WNV-infected organoids at 72 hpi). **(D)** UMAP visualisation of mock-infected and WNV-infected organoids at 24 and 72 hpi split by condition and coloured by annotated cell type. Major populations included cycling progenitors, radial glia, stressed radial glia, astroglia, intermediate progenitors, immature neurons, mature excitatory neurons, and inhibitory neurons. **(E)** Heatmap of scaled expression of representative cluster-enriched genes across unsupervised cell clusters identified in human cerebral organoids. **(F)** UMAP feature plots showing expression of canonical marker genes used for cell type annotation, including astrocyte markers (*S100B*, *SPARCL1*), neuronal markers (*RBFOX3*, *CPLX1*), and progenitor or proliferative markers (*PAX6*, *FZD2*, *MKI67*, *TOP2A*). Colour intensity indicates normalised gene expression.

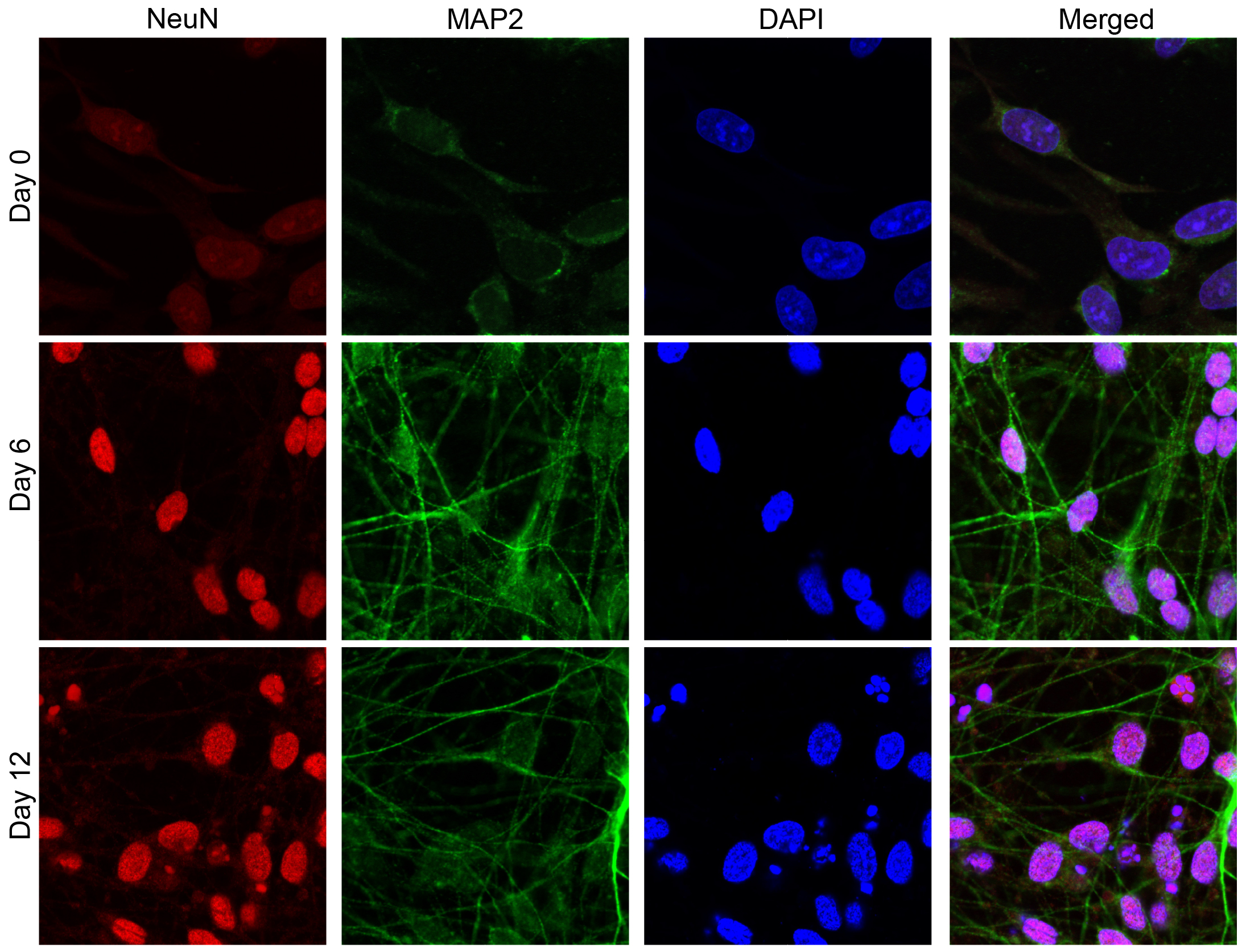

**Fig. S2. Validation of the neuronal characteristics of ReNcell VM-derived neurons.** Immunofluorescent staining of differentiating ReNcell VM cells was performed at 0, 6, and 12 days after the differentiation was induced through growth factor withdrawal. Cells were fixed and stained for pan-neuronal marker MAP2 and mature neuron marker NeuN. Nuclei were counterstained with DAPI.

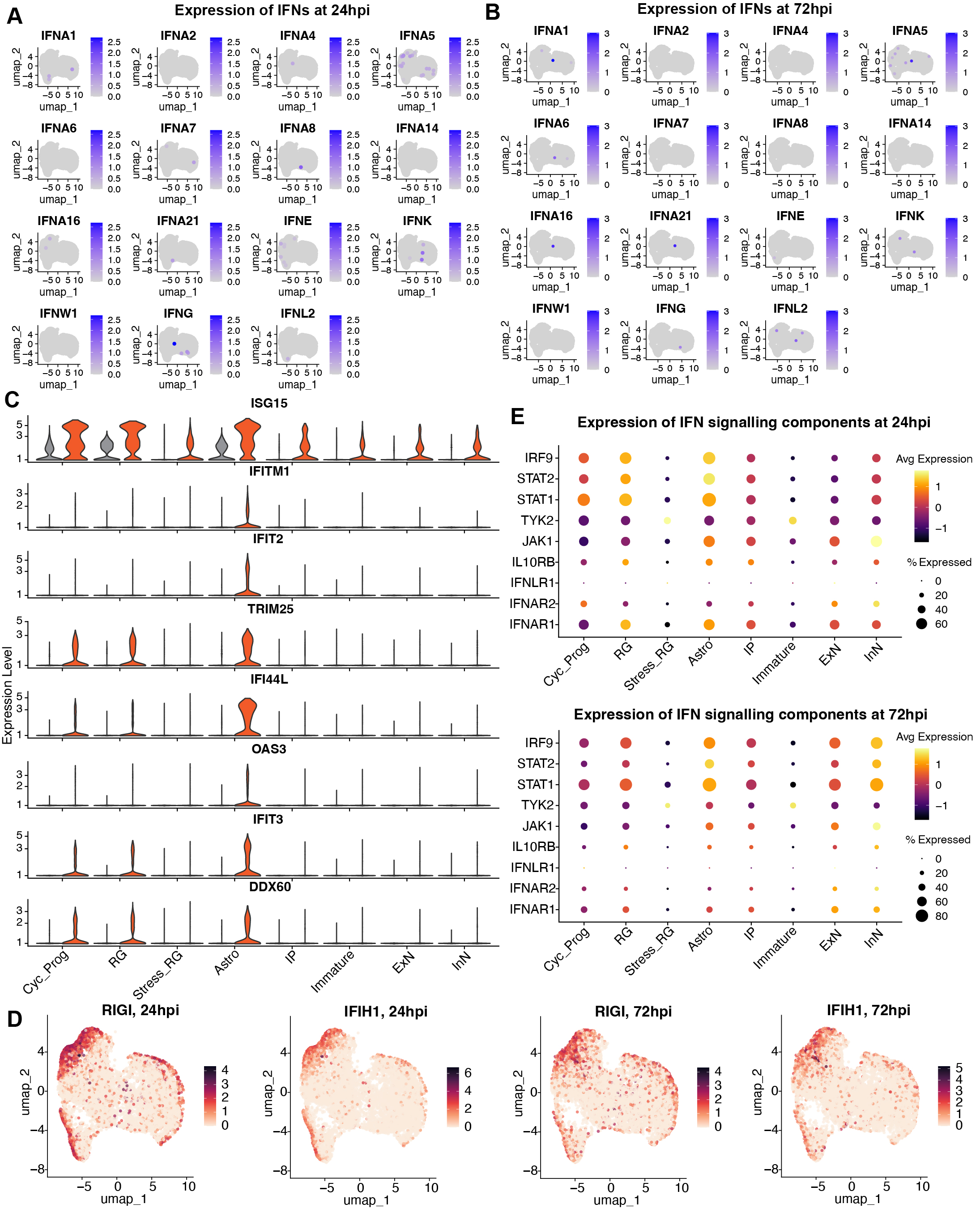

**Fig. S3. Expression of the innate immune genes in WNV-infected human cerebral organoids determined by scRNA-Seq. (A, B)** Expression of type I, type II, and type III interferon genes in WNV-infected human cerebral organoids at 24 hpi **(A)** and 72 hpi **(B)**. Each point represents a single cell positioned on the UMAP embedding, colour intensity indicates normalized gene expression. **(C)** Expression of representative interferon-stimulated genes (ISGs) across distinct cell populations in mock-infected (grey) and WNV-infected (orange) organoids at 24 hpi. **(D)** Feature plots showing expression of the viral RNA sensors MDA5 (*IFIH1*) and RIG-I (*DDX58*) in individual cells of WNV-infected cerebral organoids. **(E)** Expression of interferon signalling components across distinct cell populations in WNV-infected cerebral organoids at 24 hpi. Dot size indicates the percentage of cells expressing each gene, and colour indicates average normalized expression. Cell populations are abbreviated as cycling progenitors (Cyc_Prog), radial glia (RG), stressed radial glia (Stress_RG), astroglia (Astro), intermediate progenitors (IP), immature neurons (Immature), excitatory neurons (ExN), and inhibitory neurons (InN).

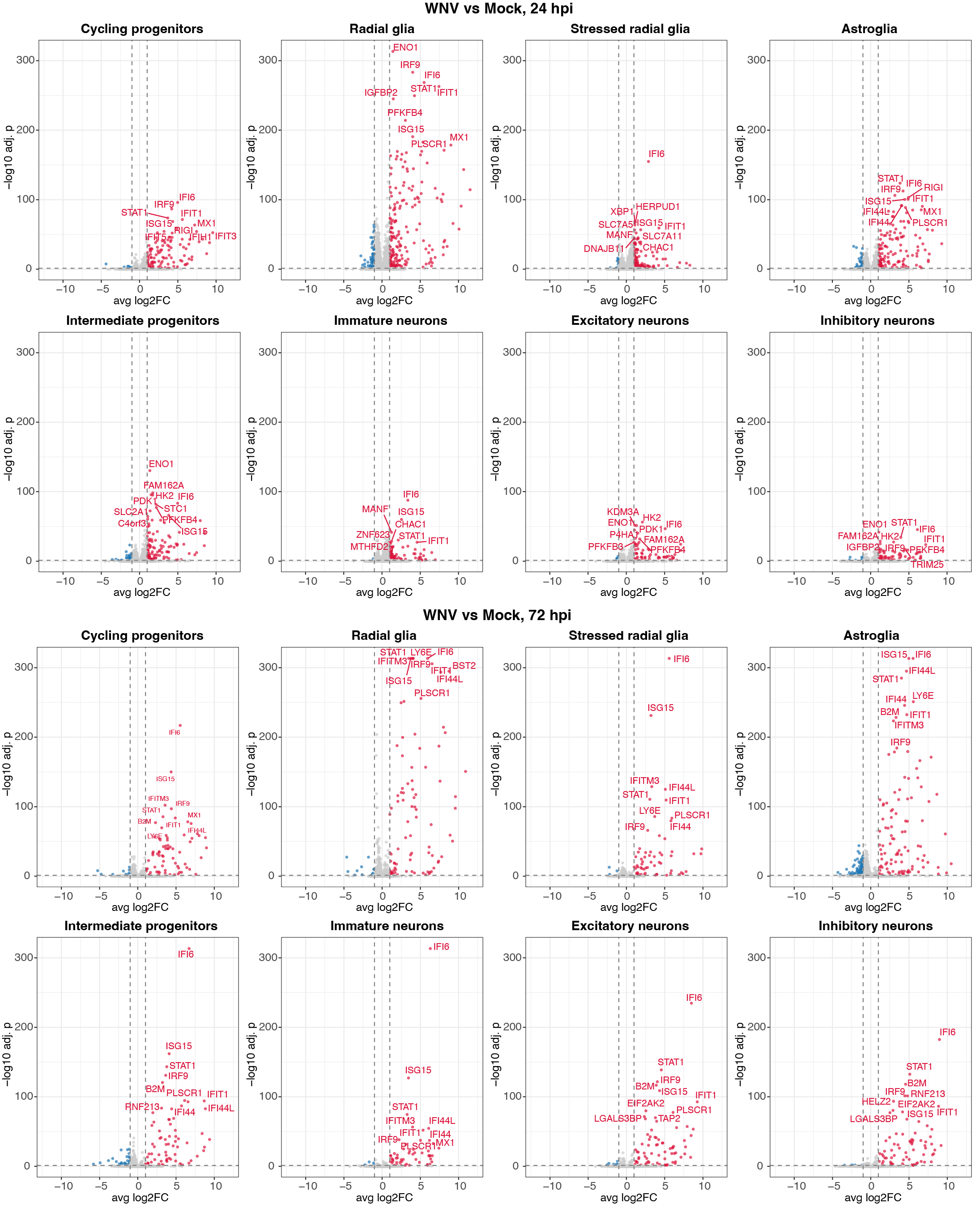

**Fig. S4. Differential gene expression in WNV-infected cerebral organoids across different cell types.** Volcano plots show differentially expressed genes (DEGs) identified in each annotated cell cluster for two comparisons: WNV at 24 hpi versus mock (top) and WNV at 72hpi versus mock (bottom). Differential expression was computed on log-normalized counts using two-sided Wilcoxon rank-sum tests, p-values were adjusted for multiple comparisons using the Bonferroni method across all genes tested. Red points indicate significantly up-regulated genes and blue points significantly down-regulated genes (adjusted P < 0.05 and log₂FC > 1). Dashed lines mark the significance thresholds. The 10 genes with the smallest adjusted P values per cluster are labelled. Axes are held constant across all panels to allow direct visual comparison.

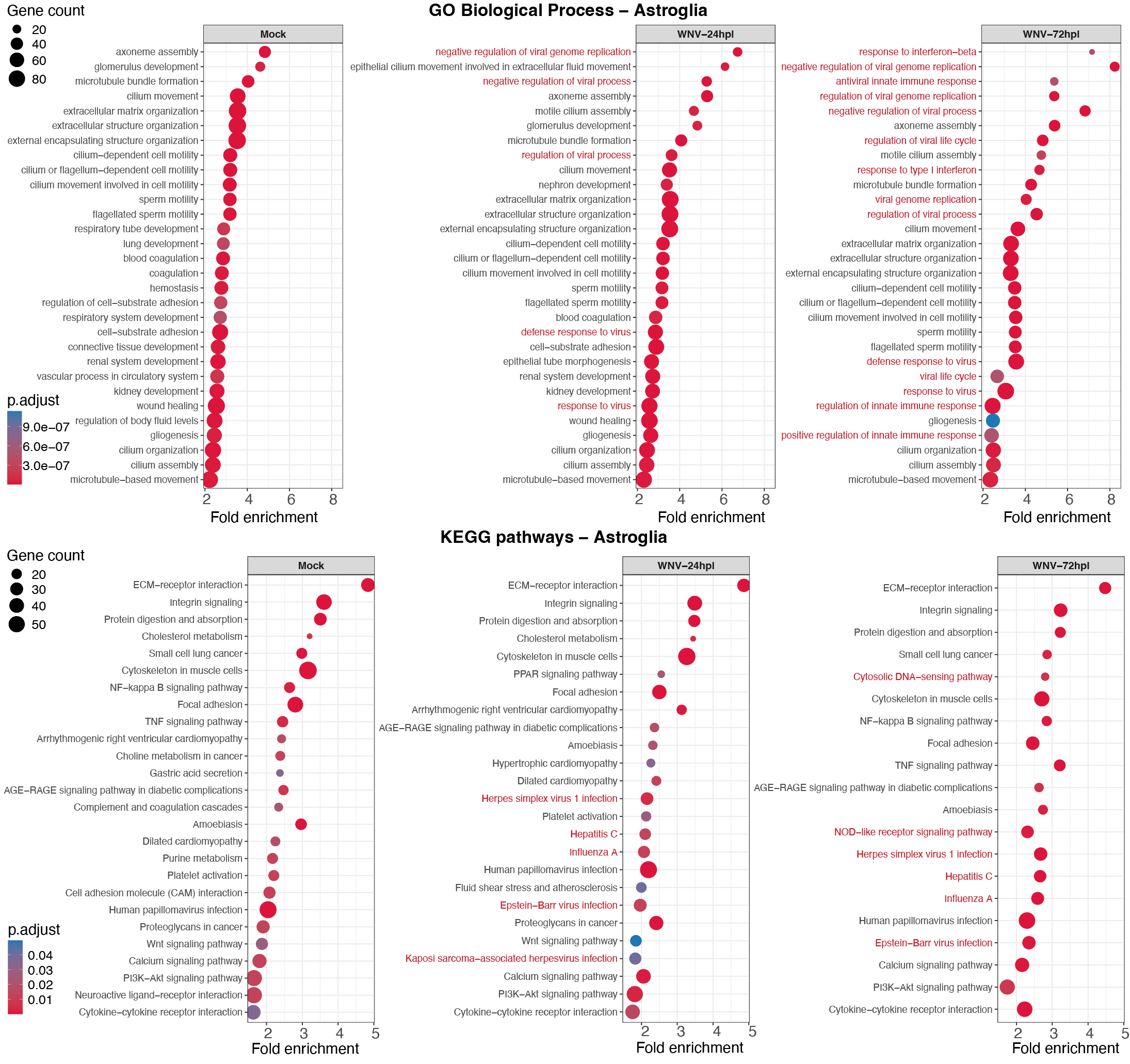

**Fig. S5. Gene Ontology terms and KEGG pathway associated with astroglia-enriched genes in WNV-infected and uninfected (Mock) human cerebral organoids.** Dot plots show enriched Gene Ontology Biological Process terms and KEGG pathways for genes that have enriched expression in astroglia relative to all other neural cell types. Enriched genes were identified by two-sided Wilcoxon rank-sum test (astroglia versus remaining clusters). Genes with Bonferroni-adjusted P < 0.05 and average log₂ fold change > 1 were submitted for over-representation analysis in clusterProfiler. Enrichment P values were calculated by one-sided hypergeometric test and adjusted for multiple comparisons by the Benjamini-Hochberg method; only terms with adjusted P < 0.05 are shown, and the 30 most significant per condition are displayed.

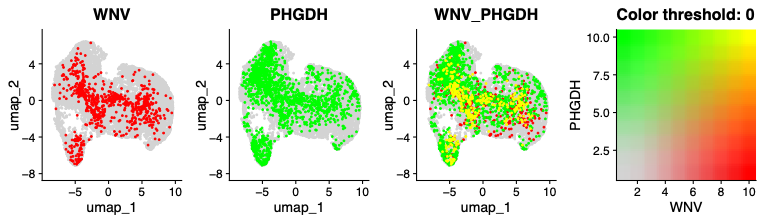

**Fig. S6. Co-detection of WNV RNA and PHGDH gene expression in infected human cerebral organoids.** Uniform manifold approximation and projection (UMAP) embeddings of single cell transcriptomes from WNV-infected cerebral organoids at 72 hpi coloured by expression of the indicated features. Left to right: WNV viral RNA alone (red), *PHGDH* alone (green), and the two features overlaid (WNV_PHGDH), with co-expressing cells appearing yellow. Expression values are log-normalized counts, each point represents one cell.
